# Dysregulated heparan sulfate proteoglycan metabolism promotes Ewing sarcoma tumor growth

**DOI:** 10.1101/2021.05.25.445683

**Authors:** Elena Vasileva, Mikako Warren, Timothy J. Triche, James F. Amatruda

## Abstract

The Ewing sarcoma family of tumors is a group of malignant small round blue cell tumors (SRBCTs) that affects children, adolescents and young adults. The tumors are characterized by reciprocal chromosomal translocations that generate chimeric fusion oncogenes, the most common of which is EWSR1-FLI1. Survival is extremely poor for patients with metastatic or relapsed disease, and no molecularly-targeted therapy for this disease currently exists. The absence of a reliable genetic animal model of Ewing sarcoma has impaired investigation of tumor cell/microenvironmental interactions *in vivo*. We have developed a new genetic model of Ewing sarcoma based on Cre-inducible expression of human *EWSR1-FLI1* in wild type zebrafish, which causes rapid onset of SRBCTs at high penetrance. The tumors express canonical EWSR1-FLI1 target genes and stain for known Ewing sarcoma markers including CD99. Growth of tumors is associated with activation of the MAPK/ERK pathway, which we link to dysregulated extracellular matrix metabolism in general and heparan sulfate catabolism in particular. Targeting heparan sulfate proteoglycans with the specific heparan sulfate antagonist Surfen reduces ERK1/2 signaling and decreases tumorigenicity of Ewing sarcoma cells *in vitro* and *in vivo*. These results highlight the important role of the extracellular matrix in Ewing sarcoma tumor growth and the potential of agents targeting proteoglycan metabolism as novel therapies for this disease.

---

Ewing sarcoma is an aggressive sarcoma of bone and soft tissue with a peak incidence in adolescents and young adults (Gaspar et al., 2015). Diagnosis relies on histologic and molecular analysis of biopsy specimens or surgically resected tumor tissue. Histologic examination of the tumor typically reveals sheets of small, round, blue cells with a prominent nucleus and scant cytoplasm. Approximately 20–25% of patients present with metastases at diagnosis that are often resistant to intensive therapy (Gaspar et al., 2015). Standard multimodal therapy for patients with small round blue cell tumor (SRBCT) includes surgical resection and/or local radiotherapy as well as intensive multi-agent chemotherapy (Grünewald et al., 2018). Ewing sarcoma family tumors are characterized by the presence of reciprocal chromosomal translocations that fuse a member of the FET family of RNA-binding proteins (encoded by FUS, EWSR1 and TAF15), with different members of the ETS (E26-specific) family of transcription factors. The most common oncofusion found in 85% of cases is *EWSR1-FLI1*(Delattre et al., 1992; Grünewald et al., 2018). These chimeric fusion oncoproteins act as aberrant transcription factors deregulating hundreds of genes (for example, genes involved in cell-cycle regulation, cell migration and proliferation) by binding DNA enriched with GGAA motifs (Gangwal et al., 2008; Guillon et al., 2009; Johnson et al., 2017). These and other studies, made on patient-derived tumor tissue and cell lines, have provided great insight into the molecular mechanisms of fusion protein function. However, in the absence of a representative *in vivo* model, other questions, including the cell of origin of Ewing sarcoma, mechanisms of tumor initiation and the relationships between tumor and host cells, remain a subject of constant debate.

ERK1 and ERK2 are key effectors in the Ras-Raf-MEK-ERK signal transduction cascade. The phosphorylation of ERK1/2 is required for the activation and subsequent phosphorylation of hundreds of cytoplasmic and nuclear substrates including transcription factors regulating cell adhesion, migration, and proliferation (Meloche and Pouysségur, 2007). ERK1/2 activity is required for transformation of NIH-3T3 cells by EWSR1-FLI1 (Silvany et al., 2000). In therapy-naive Ewing sarcoma samples, combined expression of pAKT, pmTOR, and pERK predicted worse progression-free and overall survival (Van De Luijtgaarden et al., 2013). ERK is a downstream target of Insulin-like growth factor signaling, however the prognostic impact of IGF-1 and IGF-1 receptor expression in Ewing sarcoma is controversial (Scotlandi et al., 2011). These findings raise the question of precisely which mechanisms drive ERK1/2 signaling in Ewing sarcoma, including the possible role of the tumor microenvironment.

The tumor microenvironment (TME) plays an important role in determining tumor cell growth, survival, and response to treatment. The tumor microenvironment is composed of various cell types embedded in an altered extracellular matrix (ECM). The ECM is a three-dimensional network of extracellular proteins, collagen, glycoproteins and signaling molecules that provide structural and biochemical support to surrounding cells, as well as playing an essential role in signal transduction (Kim et al., 2011). Proteoglycans are key molecular effectors of the cell surface and pericellular microenvironment with essential roles in signal transduction and regulating cell adhesion, migration and differentiation. Signaling between cells is modulated by proteoglycan activity at the cell membrane (Elfenbein and Simons, 2010). They have an ability to interact with both ligands and receptors that could directly affect cancer growth (Edwards, 2012; Iozzo and Sanderson, 2011; Multhaupt et al., 2016; Mythreye and Blobe, 2009). Importantly heparan sulfate proteoglycans play an essential role in differentiation and migration of both neural crest and mesenchymal cells – both of them are considered as potential cell of origin of Ewing sarcoma (Henderson and Copp, 1997; Long and Huttner, 2019; Papy-Garcia and Albanese, 2017; Yaylaci et al., 2016). To date, however, little is known about the role of proteoglycans and activation of ERK1/2 signaling in Ewing sarcoma development.

A genetic animal model of Ewing sarcoma family of tumors would thus be highly valuable as a complement to existing xenograft models for exploration of cooperating genetic factors for tumor development and for testing the role of the complex microenvironment in tumor progression. However, multiple attempts to create genetically-engineered mouse models of Ewing sarcoma have not yielded a tractable model, likely due to the developmental toxicity of heterotopic expression of the oncofusion (Minas et al., 2017). Previously we demonstrated that transposon-mediated expression of human *EWSR1-FLI1* drives small round blue cell tumor formation in zebrafish from 6 to 19 months of age (Leacock et al., 2012). While serving as proof of principle that *EWSR1-FLI1* is tumorigenic in fish, the model had some limitations, including low penetrance, requirement for tp53 deficiency, and a low incidence of other tumor types such as leukemias.

Here we describe Cre-inducible invasive model of Ewing Sarcoma in zebrafish that reproducibly develops tumors in wild type backgrounds. We characterize tumors and show that they recapitulate the main aspects of the human disease. Using this model, we show that tumor growth is associated with activation of the ERK1/2 signaling pathway, and that EWSR1-FLI1 upregulates expression of proteins involved in extracellular matrix reorganization and heparan sulfate proteoglycan catabolism. We demonstrate that Surfen, a heparan sulfate antagonist, effectively reduces proteoglycan-mediated activation of ERK1/2 signaling in Ewing sarcoma cell lines and zebrafish models, leading to decreased proliferation of Ewing sarcoma tumor cells *in vitro* and *in vivo*.

## Results

### Cre-inducible expression of *EWSR1-FLI1* drives SRBCT development in zebrafish

We reasoned that the low penetrance of tumors and requirement for loss of tp53 function in our original zebrafish Ewing sarcoma model (Leacock et al., 2012) might be due to developmental toxicity of *EWSR1-FLI1*, similar to what was described in mouse models (Minas et al., 2017). This is especially true since microinjection is performed into single cell-stage embryos, allowing integration of transposons into cell populations early in development. We therefore tested several strategies, beginning with expression of *EWSR1-FLI1* under ubiquitous and tissue specific promoters in wild type zebrafish embryos (Fig. 1A). All constructs incorporated eGFP linked to *EWSR1-FLI1* via a viral 2A linkage, to allow fluorescent labeling of *EWSR1-FLI1-expressing* cells via an unfused eGFP (Provost et al., 2007). Consistent with our previous data (Leacock et al., 2012), constitutive expression of *eGFP-2A-EWSR1-FLI1* under the *beta-actin* promoter in wildtype fish caused high embryonic lethality and low incidence of tumor formation. We obtained similar results with other ubiquitous promoters including *cmv* and ubiquitin (*ubi*). On the other hand, expression of *EWSR1-FLI1* from tissue-specific promoters (*sox9b*, *fli1a* and *mitfa*) failed to produce tumors (Supplemental Fig.1A). These promoters become active only during somitogenesis or at later stages, suggesting that the developmental stages or lineages marked by these promoters were no longer susceptible to transformation by *EWSR1-FLI1*. Based on these results, we suspected that at early stages of development, beyond the initial pre-gastrulation stage but prior to the onset of somitogenesis, there might be a population of cells that would be susceptible to transformation. We accordingly designed a Cre-inducible allele (loxP-DsRed-STOP-loxP-GFP-2A-*EWSR1-FLI1*, henceforth Red-STOP-Green or “*RSG*”-*EWSR1-FLI1*) that would allow more control over the timing of *EWSR1-FLI1* expression. Injection of *RSG-EWSR1-FLI1* transposon and *Tol2* transposase, along with mRNA encoding Cre, allows for a short delay in expression of *EWSR1-FLI1*, as the *Cre* mRNA must be translated before recombination can occur. Initial attempts to generate tumors with *RSG*-*EWSR1-FLI1* driven by the beta-actin promoter (Supplemental Fig. 1B, 2B,C) caused early apoptosis (Supplemental Fig. 2A) and the growth of cranial cell masses (Supplemental Fig. 2D), however failed to recapitulate the histologic appearance of Ewing sarcoma (Supplemental Fig. 2E).

**Figure 1.**
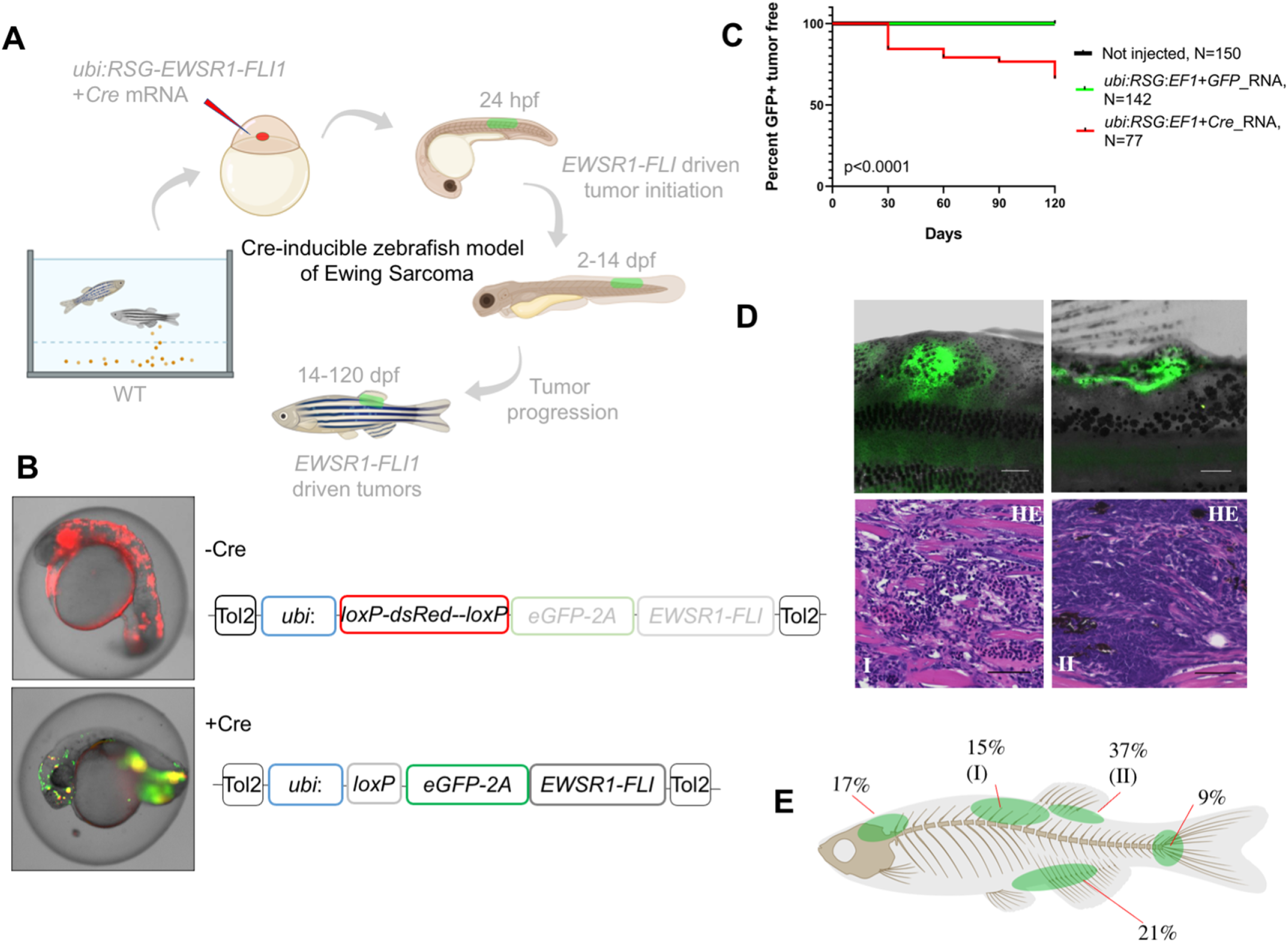
Cre-inducible expression of *EWSR1-FLI1* drives SRBCT development in zebrafish. **A,** Overview of experimental method. The Tol2 transposon system was used to integrate human *EWSR1-FLI1* into the wild type zebrafish genome by microinjection into singlecell stage embryos. GFP-positive fish were monitored up to 4 months. **B**, **C**onstructs for Cre-inducible expression of *EWSR1-FLI1*. **C**, Incidence of GFP + tumors detected in *Ubi*:*RSG-EWSR1-FLI1*; *Cre* RNA (n = 77) injected zebrafish versus *Ubi*:*RSG-EWSR1-FLI1*; *GFP* RNA (n = 142) and uninjected controls (N=150) in a wildtype genetic background. (p<0.001 by log-rank test). **D**, Representative images of GFP-positive tumors in zebrafish (top panel) and H&E staining of tumor sections (bottom panel): (I) SRBCT with diffuse skeletal muscle infiltration, (II) visceral SRBCT arising from fin dorsal radial bone. Scale bars, 100 μm. **E**, Percent tumor incidence at different anatomic sites.

In contrast, co-injections of *Cre* mRNA with expression constructs in which *RSG-EWSR1-FLI1* is driven by the ubiquitin promoter led to robust, mosaic expression of *EWSR1-FLI1* and *GFP* (Fig.1 A, B). To estimate the frequency of tumor development using this approach, injected fish were sorted for GFP at 14 days post fertilization (dpf) and a cohort of 77 fish was monitored up to 4 months. To control for promoter leakiness, we injected embryos with *ubi*:*RSG-EWSR1-FLI1* in the presence of GFP RNA (142 fish). Uninjected embryos were used as an additional negative control (150 fish). We found that 34% of GFP positive fish developed tumors (Fig. 1C, Supplemental Table 1) exhibiting SRBCT histology (Fig. 1D, Supplemental Fig. 3C). No tumors were observed in either control group. Among the fish developing tumors, 42% developed a single tumor, while 58% of fish had 2 or more tumors (Supplemental Fig. 3A, Table 1). *EWSR1-FLI1* driven tumors arose most frequently in the dorsal regions of the fish (Fig. 1E). 37% of tumors were SRBCTs associated with the dorsal fin radial bones, while a further 15% of tumors arising from similar locations showed deep muscle invasiveness and were characterized as SRBCT with diffuse striated muscle infiltration. 9% and 21% of tumors were associated with the skeleton of caudal and anal fins, respectively. Finally, we observed several tumors that appeared to arise from the cranial skeleton or the brain. In summary, the inducible *ubi*:*GFP2A-EWSR1-FLI1* allele efficiently induced bone and soft-tissue small round blue cell sarcomas in fish, without requirement for impaired tp53 function.

### *EWSR1-FLI1* driven tumors in zebrafish recapitulate the main aspects of human Ewing sarcoma

To further characterize the zebrafish model, we performed immunohistochemistry (IHC) on tumors with antibodies specific for human FLI1 and CD99 (Fig. 2A, Supplemental Fig. 4). As expected, tumor cells had nuclear localization of EWSR1-FLI1, whereas control fish had no expression of FLI1 in the corresponding sites (Supplemental Fig. 4 top panel). CD99 is a cell surface glycoprotein that serves as a sensitive, clinically useful marker for Ewing sarcoma (Muhammad et al., 2012). IHC confirmed the presence of CD99 on the cell surface of zebrafish Ewing sarcoma tumor cells (Fig. 2A). Ewing sarcoma cells are characterized by presence of the glycogen and are positive for Periodic acid-Schiff (PAS) staining (Muhammad et al., 2012).

**Figure 2.**
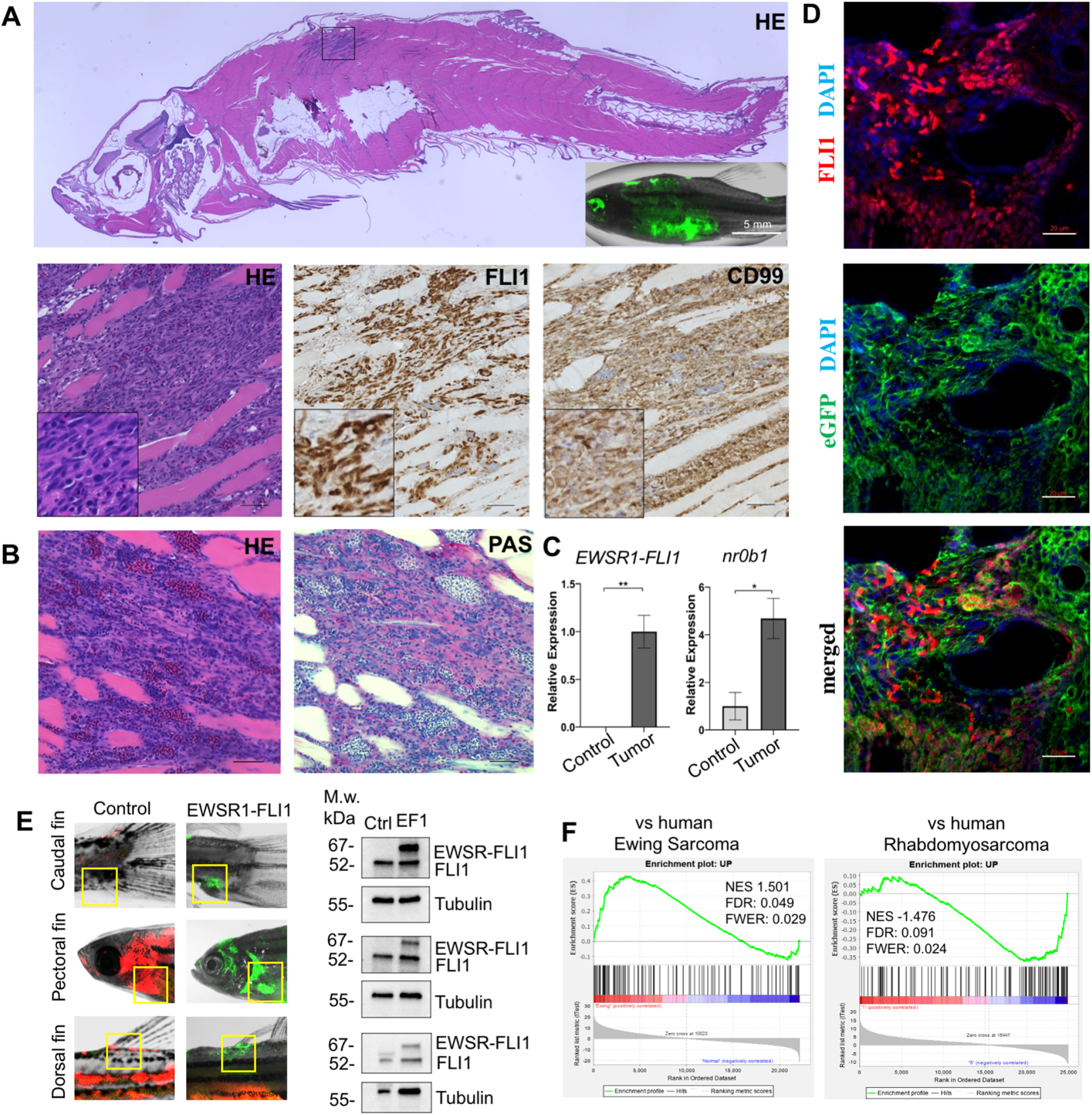
Zebrafish tumors phenocopy human Ewing sarcoma. **A,** Representative image of Ewing sarcoma tumor in WT zebrafish. H&E staining, IHC staining with anti-FLI1 and anti-CD99 antibodies. Scale bars, 100 μm. **B,** PAS staining of zebrafish tumors. Scale bars, 50 μm. **C,** Relative expression of human *EWSR1-FLI* and zebrafish *nr0b1* in normal and tumor tissues. Error bars represent SEM, N = 3, * indicates p<0.05, ** indicates p<0.01, two-tailed Student’s t-test. **D,** Immunofluorescence staining of zebrafish tumor with anti-GFP and anti-FLI1 antibodies. Scale bars, 20μm. **E,** Validation of *EWSR1-FLI1* expression in tumors at different sites by immunoblotting. **F,** GSEA comparing the enrichment of common upregulated proteins at dorsal, caudal and pectoral fin tumors to human Ewing sarcoma (GSE17674) and human Rhabdomyosarcoma (GSE108022) datasets.

Consistently, zebrafish tumors were positive for PAS staining (Fig. 2B). *NR0B1* is a target of EWSR1-FLI1 that directly modulates transcription and oncogenesis in Ewing sarcoma (Kelsey C. Martin Mhatre V. Ho, 2009). Analysis of relative RNA expression showed upregulated expression of *nr0b1* in tumor tissues compared to normal (Fig. 2C), correlating with *EWSR1-FLI1* expression level in tumor samples. Recently, it was shown that Ewing sarcoma cells exhibit cell to cell heterogeneity affecting proliferative and migratory potential of tumor cells (Franzetti et al., 2017). To test whether zebrafish tumor cells express *EWSR1-FLI1* on different levels we performed immunofluorescence staining of tumor sections with anti-GFP and anti-FLI1 antibodies (Fig.2 D). GFP staining was used to label all tumor cells within the tumor, while FLI1 staining was implemented to detect the EWSR1-FLI oncofusion in tumor cells (Fig.2D). Staining of zebrafish tumors showed that cells have varying levels of EWSR1-FLI1. Thus, the zebrafish model appears to recapitulate the cell-to cell heterogeneity found in human Ewing Sarcoma cells.

To test whether tumors express EWSR1-FLI1 on the protein level we collected tumor tissues at different sites (dorsal, caudal and pectoral fins) and processed them for immunoblot analysis. All tumor samples expressed EWSR1-FLI1 (Fig. 2E). We next performed LC-MS/MS mass spectrometry analysis of protein samples made from tumors located on dorsal fin, caudal fin and pectoral fin. Normal tissues from the corresponding sites were used as controls. We first identified a set of 194 proteins that were commonly upregulated in the zebrafish tumors compared to control tissues (pval<0.05). To test the similarity of fish and human Ewing sarcoma, we used gene set enrichment analysis (GSEA) (Subramanian et al., 2005) to test the enrichment of the 194 genes in a publicly available RNA-seq dataset of 44 human Ewing sarcoma tumors and 18 normal samples Series (GSE17674). As a further test of specificity, we compared the fish tumors to another RNA-Seq dataset of 44 human rhabdomyosarcoma tumors and 5 normal samples (GSE108022). GSEA showed enrichment of zebrafish tumor upregulated genes (NES = 1.501, FWER=0.029) in the human Ewing Sarcoma dataset but not in human rhabdomyosarcoma (NES = −1.476, FWER=0.024). Taken together, the histologic appearance, marker expression and gene expression establish the similarity of zebrafish EWSR1-FLI1-induced sarcomas to human Ewing sarcoma.

### Characterization of embryonic model of ES

To characterize the effects of *EWSR1-FLI1* expression during early stages of zebrafish development we integrated the *ubi*:*RSG-EWSR1-FLI1* cassette into the zebrafish genome in the presence of *Cre* mRNA (Fig. 3A top panel). Negative controls included co-injection of *Cre* mRNA with an *ubi*:*RSG* construct that lacks *EWSR1-FLI1* (Fig. 3A middle panel) and coinjection of *ubi*:*RSG-EWSR1-FLI* with *GFP* mRNA (Fig. 3A bottom panel). Embryos were imaged at 12 hours post-fertilization (hpf), 24 hpf and 5 dpf. The timeline demonstrates that GFP driven by the *ubi* promoter is expressed broadly, including throughout the muscle (Fig. 3A, middle panel). However, zebrafish expressing *eGFP-2A-EWSR1-FLI* under the *ubi* promoter have a distinct distribution pattern of GFP-positive cells (Fig. 3A upper panel), predominantly on the fish dorsum, tail and fins (Fig. 3A upper panel).

**Figure 3.**
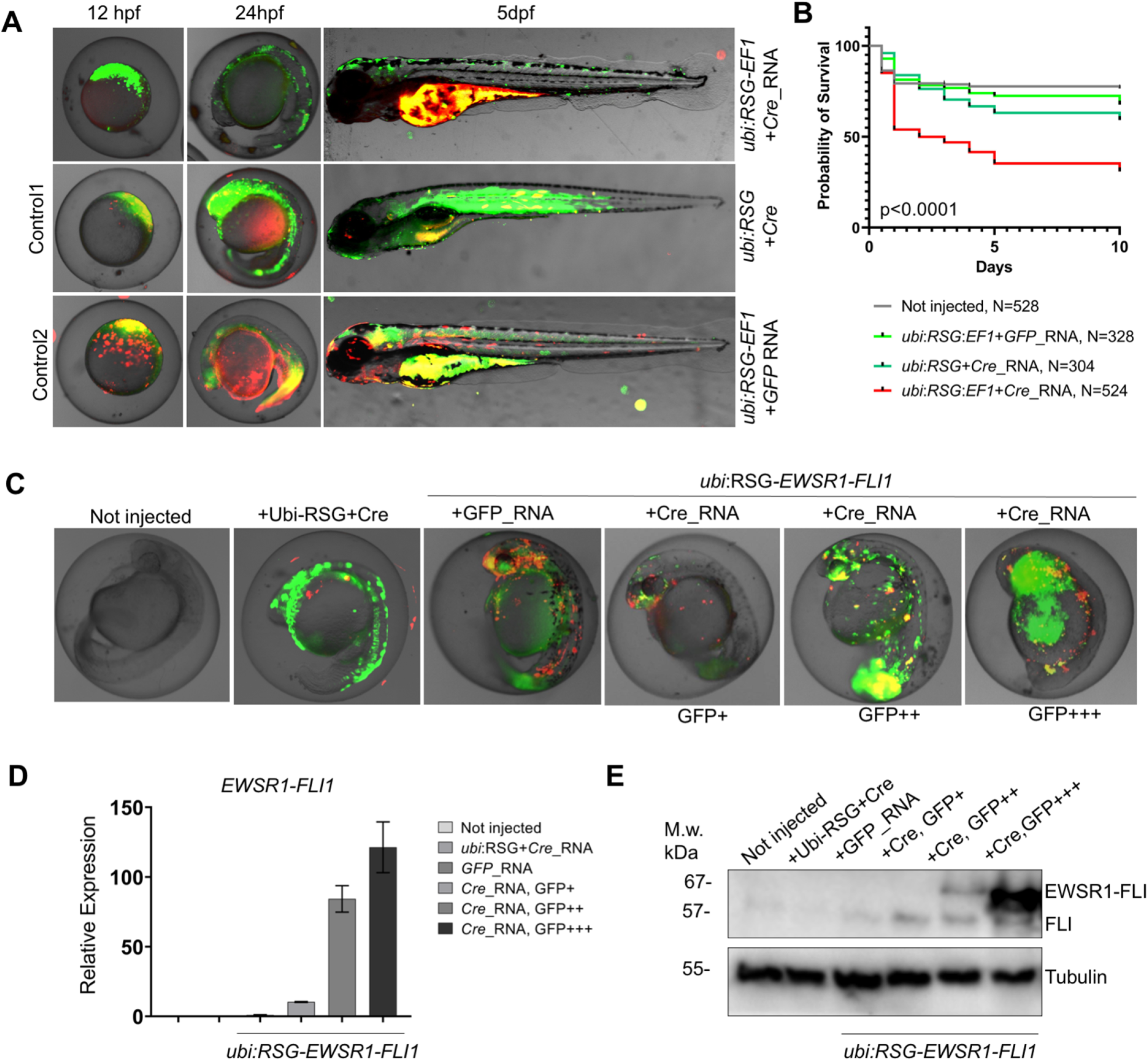
Validation of *EWSR1-FLI* expression in zebrafish embryos. **A,** Timeline of zebrafish development after injection with *ubi*:*RSG-EWSR1-FL* plus *Cre*_mRNA (top panel). Zebrafish injected with *ubi*:*RSG-EWSR1-FLI1* plus *GFP* mRNA (middle panel) or *ubi*:RSG plus *Cre* mRNA (bottom panel) were used as negative controls. Images were taken at 12hpf, 24hpf and 5dpf time points. **B,** Survival rate of embryos expressing *EWSR1-FLI1* during the first 10 days of development (N=524). Uninjected embryos (N=528) as well as embryos injected with *ubi*:*RSG-EWSR1-FLI1* plus *GFP* mRNA (N=328) or *ubi*:RSG plus *Cre* mRNA (N=304) were used as negative controls (p<0.001 by log-rank test). **C,** Representative image of embryos with low (GFP+), medium (GFP++) and high (GFP+++) levels of *EWSR1-FLI1* expression. **D,** Relative mRNA expression of *EWSR1-FLI1* in embryos with low (GFP+), medium (GFP++) and high (GFP+++) levels of *EWSR1-FLI1* according to GFP signal. **E,** Immunoblot confirming the expression of EWSR1-FLI1 protein in embryos with low (GFP+), medium (GFP++) and high (GFP+++) levels of EWSR1-FLI1 expression according to GFP signal.

In most murine models, *EWSR1-FLI* expression is toxic and causes embryonic lethality (Minas et al., 2017). To estimate the developmental toxicity of *EWSR1-FLI1* expression in zebrafish we performed survival analysis in a group of 508 animals. Uninjected embryos (N=528), embryos injected with *ubi*:*RSG-EWSR1-FLI1* and *GFP* mRNA (N=318) and embryos injected with *ubi*:*RSG* and *Cre* mRNA (N=298) were used as negative controls. Expression of *EWSR1-FLI1* significantly increased embryonic mortality in zebrafish (Fig. 3B).

To confirm the expression of *EWSR1-FLI1* in the zebrafish embryo model, we collected embryos expressing low (GFP+), medium (GFP++) and high (GFP+++) levels of *EWSR1-FLI1* (Fig. 3C) for analysis via qRT-PCR (Fig. 3D) or immunoblot (Fig. 3E). Upon Cre-mediated recombination, *EWSR1-FLI* is efficiently expressed in embryos on both the RNA and protein levels (Fig. 3 D,E).

In summary, the *Cre*-inducible allele leads to robust and reproducible expression of human *EWSR1-FLI1* with limited spatial distribution during early zebrafish embryogenesis. Developmental toxicity, while much less than that caused by *EWSR1-FLI* expression under *b-actin* and *cmv* promoters, does impact the survival of larvae over the first 10 days of life.

### EWSR1-FLI1 expression leads to activation of ERK1/2 signaling in zebrafish embryos and adult tumors

Our finding that expression of *EWSR1-FLI1* in early embryos leads to a high prevalence of tumors in adult zebrafish led us to examine the behavior of cells expressing *EWSR1-FLI1* in developing embryos, to understand early events in tumorigenesis. Zebrafish embryos mosaically expressing *EWSR1-FLI1*, but not control embryos, developed outgrowths visible as discrete cell masses. Immunostaining with anti-phospho-histone H3 (pH3) showed increased cell proliferation in these areas, associated with expression of *EWSR1-FLI1* as indicated by the associated GFP marker (Fig. 4A). The Ras-MAPK-ERK signaling pathway transduces signals downstream of growth factor receptors, and is an important mediator of cell proliferation during embryonic development and in cancer (Kamiya et al., 2015; Maekawa et al., 2007; Steinmetz et al., 2004; Wong et al., 2019; Yang et al., 2021; Zhou et al., 2019). To assess the contribution of MAPK-ERK signaling to formation of outgrowths, we performed immunofluorescence staining for the active, phosphorylated form of ERK1/2 (pERK1/2). Cells expressing *EWSR1-FLI1* were positive for pERK1/2 (Fig. 4B). While most pERK1/2-positive cells also expressed *EWSR1-FLI1* (Fig. 4C), some surrounding cells lacking the transgene were also marked with pH3 and pERK1/2 (Supplemental Figure 5A). Thus, activation of ERK1/2 signaling is an early event in *EWSR1-FLI* driven aberrations. To test whether the activation of ERK1/2 signaling could be found in mature zebrafish tumors, we prepared histologic sections of tumors and performed immunohistochemistry for pERK1/2 (Fig. 4D, Supplemental Figure 5B), complemented by immunoblot analysis (Fig. 4E,F). Similar to *EWSR1-FLI1* driven outgrowth in larvae, ERK1/2 is active in advanced zebrafish tumors. ERK1/2 signaling activity in tumors was heterogeneous, with focal areas showing more intense signaling activity.

**Figure 4.**
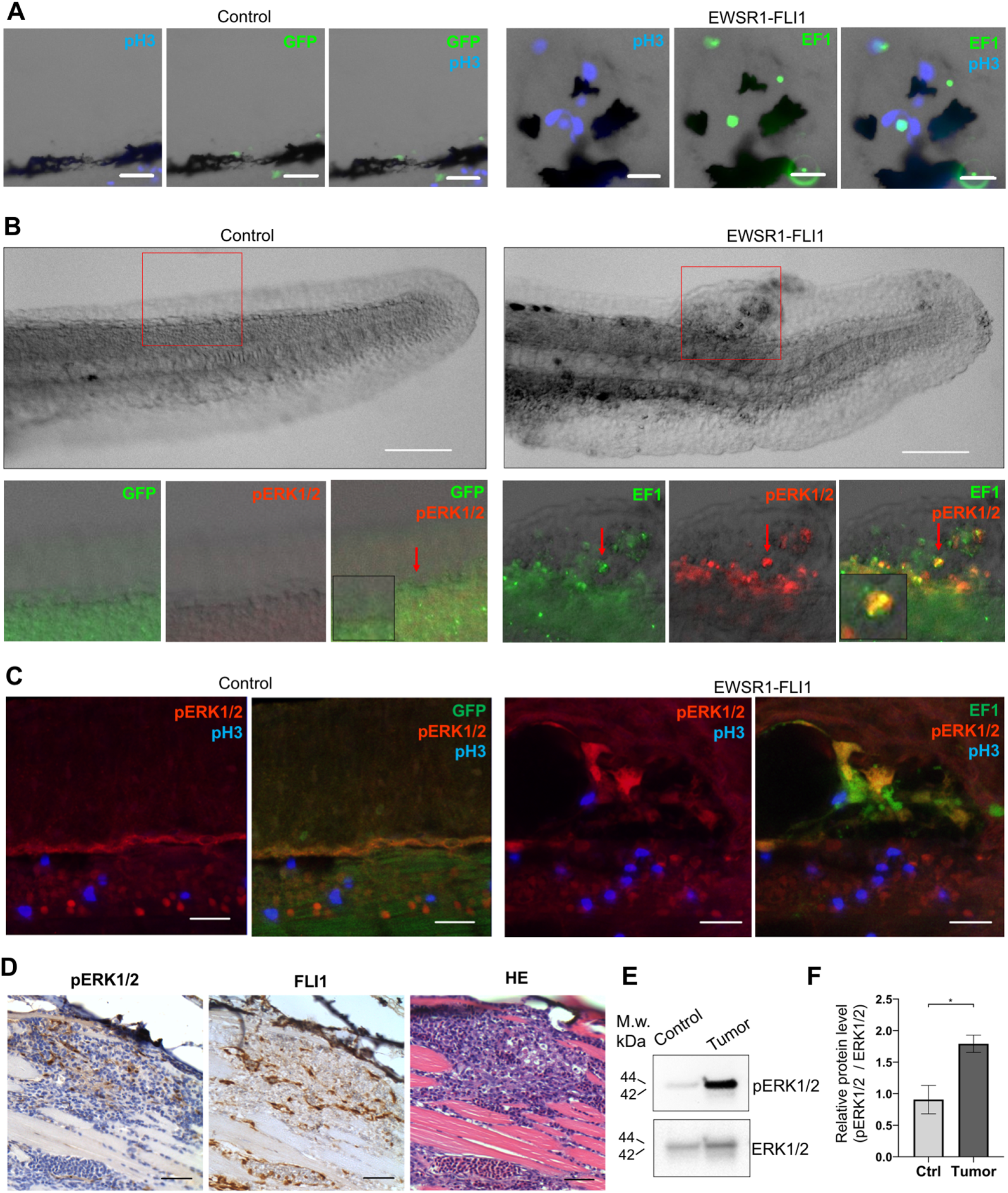
*EWSR1-FLI* expression activates the ERK1/2 signaling pathway *in vivo*. **A,** Immunofluorescent staining of *GFP* (control) and *GFP2A-EWSR1-FLI1* expressing embryos at 48hpf. Blue: phosphohistone H3. Green: GFP. A small region of the dorsal surface is shown. Scale bars, 20μm. **B,** Immunofluorescent staining of *GFP* (control) and *GFP2A-EWSR1-FLI1* expressing embryos at 24hpf for pERK1/2 (red) and GFP. Scale bars, 100μm. **C,** Immunofluorescent staining of *GFP* (control) and *GFP2A-EWSR1-FLI1* expressing embryos at 48 hpf for pERK1/2 (Red), pH3 (blue) or GFP (green). Scale bars, 20μm. **D,** Immunostaining of zebrafish tumor for pERK1/2. **E,** Immunoblot analysis of pERK1/2 and ERK1/2 levels in tumor and normal tissue. **F,** Quantification of immunoblot Error bars represent SEM, N=3, * indicates p<0.05, two-tailed Student’s t-test.

### EWSR1-FLI1 affects proteoglycan metabolism in zebrafish embryonic model

To gain further insight into the impact of *EWSR1-FLI1* expression during early tissue and organ development and the mechanisms driving increased growth factor signaling and cell proliferation, we performed mass spectrometry analysis. As above, we integrated the *ubi*:*RSG-EWSR1-FLI1* cassette into the zebrafish genome in the presence of *Cre* mRNA. The control group of embryos was co-injected with an *ubi*:*RSG* transposon and Cre mRNA. Embryos were sorted for GFP expression at 24hpf and 48hpf time points and used to generate samples for LC-MS/MS analysis. EWSR1-FLI1 significantly affects protein expression in zebrafish embryos (Fig. 5A). We identified 248 differentially expressed proteins affected by EWSR1-FLI1 at 24hpf, and 1102 at 48 hpf (Fig. 5B). Downregulated proteins comprised 65% and 73% of differentially expressed proteins at 24hpf and 48hpf, respectively, suggesting that EWSR1-FLI1 was acting mostly as a transcriptional repressor rather than an activator (Fig. 5B).

**Figure 5.**
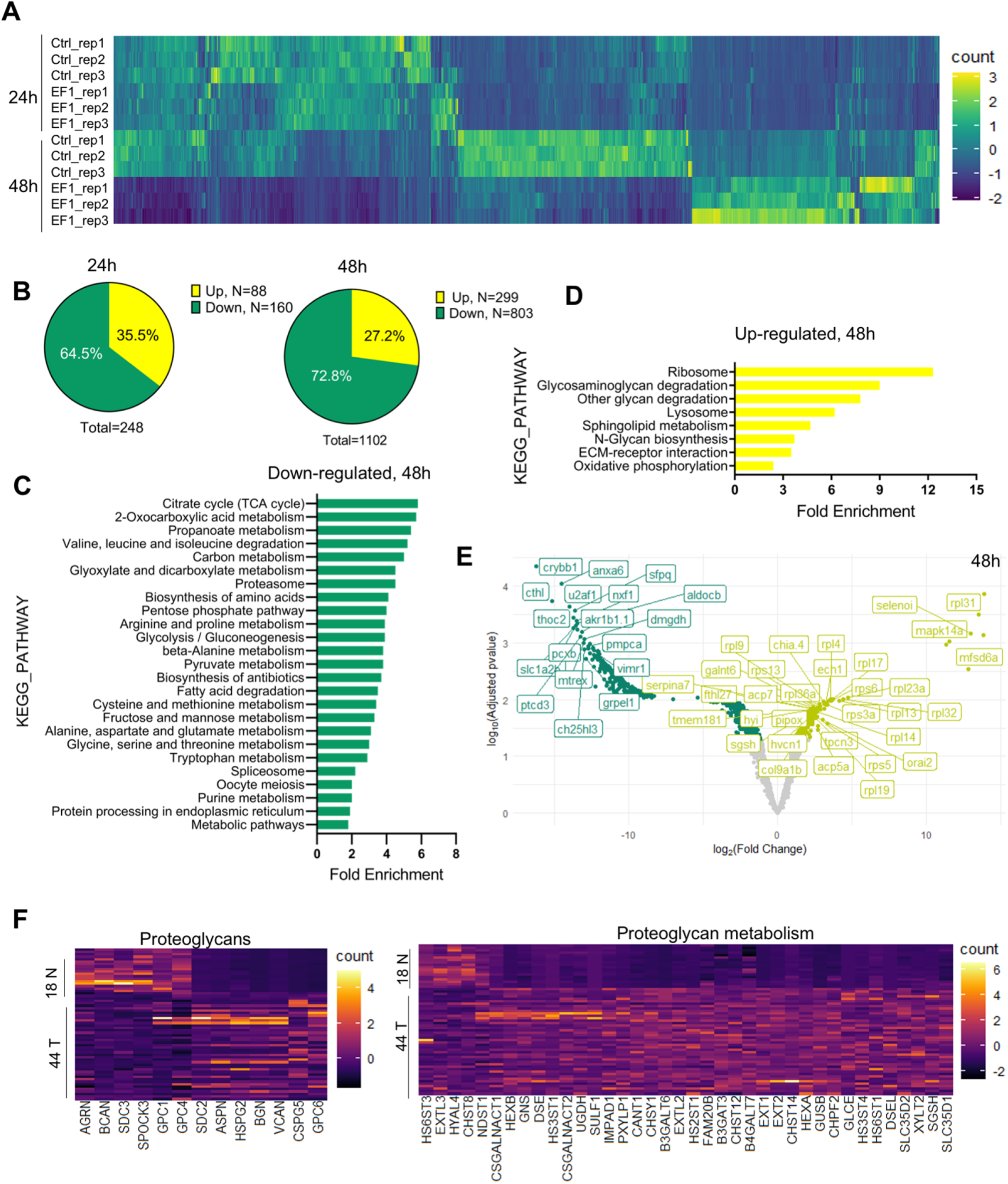
LC-MS/MS analysis of proteins affected by *EWSR1-FLI1* expression in developing zebrafish. **A,** Heat map representing the differentially expressed proteins dysregulated by *EWSR1-FLI1* at 24hpf and 48hpf. **B,** Quantitative analysis of the differentially expressed proteins at 24hpf and 48hpf. **C, D,** Gene ontology (GO) analysis of differentially expressed proteins at 48hpf. **E,** Volcano plot of significantly downregulated and upregulated proteins in *EWSR1-FLI1* expressing embryos. **F,** Heatmap of the most differentially expressed proteins involved in proteoglycan metabolism in human Ewing sarcoma compared to normal tissue (GSE17674).

Gene Ontology (GO) enrichment analysis of proteomics data at the 48hpf time point showed that downregulated proteins were involved in regulation of cell metabolism (Fig. 5C, Supplemental Table 2). Importantly, we found a strong upregulation of proteins involved in Extracellular Matrix (ECM) reorganization, proteoglycan metabolism, and protein synthesis (Fig. 5D, Supplemental Table 2). The proteomics data revealed that *EWSR1-FLI1* expression in zebrafish embryos is associated with upregulated expression of collagens *col1a1b*, *col1a2*, *col2a1a*, *col9a1b*, *col9a2 and col28a2a* (Supplemental Table 2), which are involved in ECM organization and skeletal system development. Proteins involved in proteoglycan catabolism were also significantly differentially expressed in embryos expressing *EWSR1-FLI1* Fig.5E, Supplemental Table 2). Among the top identified hits was N-sulfoglucosamine sulfohydrolase (*sgsh*), the enzyme involved in heparan sulfate proteoglycan catabolism (Fig. 5E). We also found upregulated *gnsb* and *gnsa* (N-acetylglucosamine-6-sulfatase) enzymes involved in hydrolysis of Heparan sulfate chains (Supplemental Table 2).

To compare results obtained from proteomics data of Ewing sarcoma zebrafish model with those from human Ewing sarcoma, we analyzed the profile of proteins associated with proteoglycan metabolism from microarray data of 44 Ewing sarcoma samples and 18 normal tissue samples. Expression of enzymes involved in proteoglycan metabolism was strongly upregulated in tumor samples compared to normal tissue (Fig. 5F). Specifically, we found upregulation of enzymes involved in the catabolism of heparan sulfate proteoglycans (including *SGSH*, *GNS*, *HS3ST4*, *HS3ST1*, *HS2ST1* and *HS6ST1*) in human Ewing sarcoma (Fig. 5F). Furthermore, proteoglycan metabolism was also strongly dysregulated in Ewing sarcoma tissues.

Taken together, our genetic model of Ewing Sarcoma revealed that EWSR1-FLI1 expression dysregulates normal protein expression starting from early zebrafish development. The expression of EWSR1-FLI1 is associated with strong upregulation of ECM proteins and, most notably, enzymes involved in heparan sulfate proteoglycan catabolism. Supporting these data, we show that the genes involved in proteoglycan metabolism are strongly upregulated in human Ewing sarcoma tumors.

### Surfen inhibits proteoglycan-mediated activation of ERK1/2

Proteoglycans play a key regulatory role in the interactions of cells with ECM proteins, growth factors and cytokines, and thus have major influence on cell signaling pathways that directly affect cancer growth (Edwards, 2012; Iozzo and Sanderson, 2011; Multhaupt et al., 2016; Mythreye and Blobe, 2009). Our finding that *EWSR1-FLI* expression is associated with altered proteoglycan expression and metabolism suggests that dysregulation of proteoglycans may contribute significantly to growth promoting signaling pathways in Ewing sarcoma, including ERK signaling. Thus, proteoglycan metabolism could serve as a novel target for Ewing sarcoma. To test these possibilities, we performed treatment of Ewing sarcoma cell lines with the small molecule surfen (*bis*-2-methyl-4-amino-quinolyl-6-carbamide). Surfen is a sulfated heparan sulfate antagonist with a high affinity to all sulfated GAGs (Schuksz et al., 2008). Surfen blocks sulfation and degradation of GAG chains *in vitro* and affects growth factor binding and proteoglycan-mediated signal transduction through certain cell surface growth factor receptors (Schuksz et al., 2008).

First, we tested whether surfen can block proteoglycan-mediated activation of ERK1/2 signaling in the TC32 and EWS502 Ewing sarcoma cell lines. To reduce the basal level of ERK1/2 phosphorylation we performed pre-treatment of cells with serum-free media for 4 hours, and then replaced serum-free media with the full-media in the presence of 1% DMSO or 2.5 μM, 5 μM or 10 μM surfen. After 30 minutes of treatment, cell lysates were prepared and analyzed by immunoblotting. Samples were normalized to the value of total ERK1/2. As expected, the introduction of serum led to the activation of ERK1/2 signaling (Fig. 6 A,B). Addition of 5 μM or 10 μM surfen strongly significantly inhibited ERK1/2 phosphorylation (Fig. 6A, B). Thus, blocking proteoglycan metabolism impairs ERK1/2 signaling in Ewing sarcoma cells.

**Figure 6.**
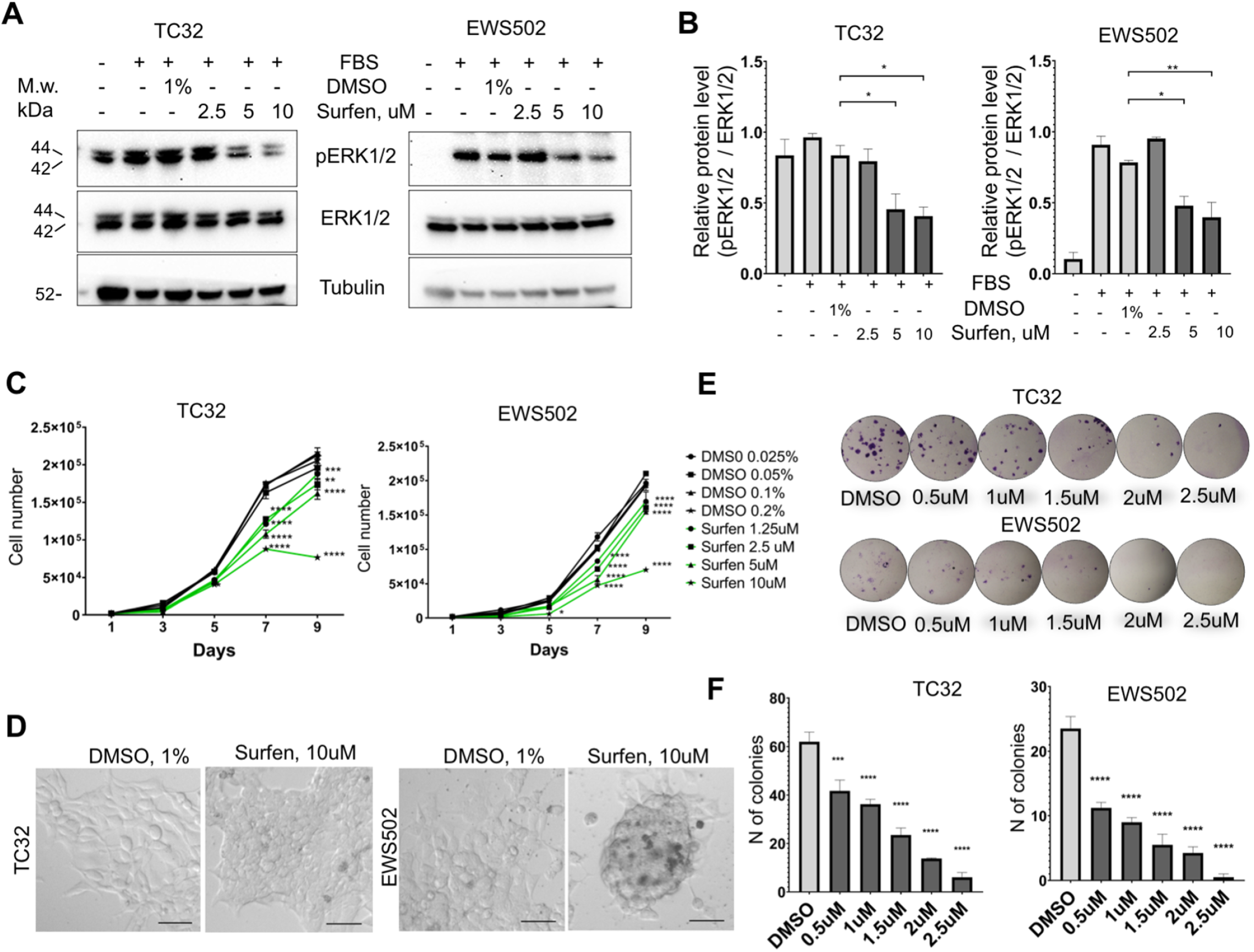
Surfen treatment impairs ERK1/2 signaling and growth of Ewing sarcoma cells. **A,** Immunoblot analysis of pERK1/2, ERK1/2 and tubulin levels in TC32 and EWS502 cells treated with surfen or DMSO. **B,** Immunoblot quantification of pERK1/2 expression level relative to ERK1/2 in TC32 and EWS502 cells treated with surfen or DMSO vehicle control. Error bars represent SEM, N=3, * indicates p<0.05, ** indicates p<0.01, based on one-way ANOVA test. **C,** Proliferation rates of TC32 and EWS502 cells treated with surfen or DMSO. Error bars represent SEM, N=3, ** indicates p<0.01, *** indicates p<0.001, **** indicates p<0.0001 based on one-way ANOVA test. **D,** Morphological changes in TC32 and EWS502 cells after surfen or DMSO treatment. **D,** Clonogenic assay of TC32 and EWS502 cells treated with Surfen or DMSO. Error bars represent SEM, N=4, *** indicates p<0.001, **** indicates p<0.0001 based on one-way ANOVA test.

We next tested the effect of surfen treatment on the proliferation rates of TC32 and EWS502 cells exposed to surfen or DMSO vehicle. Treatment with surfen significantly reduces the proliferation rates of Ewing sarcoma cells (Fig. 6C). Importantly, treatment of cells with surfen at 10 μM resulted in a decrease of TC32 proliferation by 62.7%, and EWS502 by 63.4% compared to control cells treated with 0.2% DMSO (Fig. 6C). Surfen treatment caused morphological changes in TC32 and EWS502 cells (Fig. 6D), consistent with predicted effects on cell adhesion.

To estimate cell survival and the ability of a single cell to grow into a colony we performed a clonogenic assay under low-adhesive conditions. We seeded 500 cells/well in a 24 well low adhesive plate treated with surfen or 1% DMSO, under serum-deprived conditions. Serum was added 12 hours later to stimulate growth factor-mediated signaling. The colony number was analyzed after 2 or 3 weeks of incubation for TC32 and EWS502, respectively. Treatment of cells with surfen led to a significant reduction in colony number for both cell lines (Fig. 6E, F). More specifically, the treatment of TC32 cell with 2.5 μM surfen led to a 90.3% decrease of colony number while treatment of EWS502 cells with 2.5 μM surfen resulted in a 97.8% decrease of colony forming units. Thus, surfen targets the heparan sulfate proteoglycan-mediated activation of ERK1/2 signaling in Ewing sarcoma cells, inhibiting cancer cell proliferation and cell survival.

### Surfen inhibits EWSR1-FLI1 mediated growth in zebrafish model

We next tested whether surfen could inhibit the formation of *EWSR1-FLI* driven outgrowths *in vivo* in our zebrafish model. As described above, we generated fish mosaically expressing *EWSR1-FLI1*. Fish with GFP-positive outgrowths were identified at 24hpf and treated with at 2 μM surfen or 0.2% DMSO. Fish were imaged after 24 and 48 hours of treatment (Fig. 7A). Consistent with results on human cells, surfen inhibited outgrowth development in the zebrafish embryo model of Ewing sarcoma.

**Figure 7.**
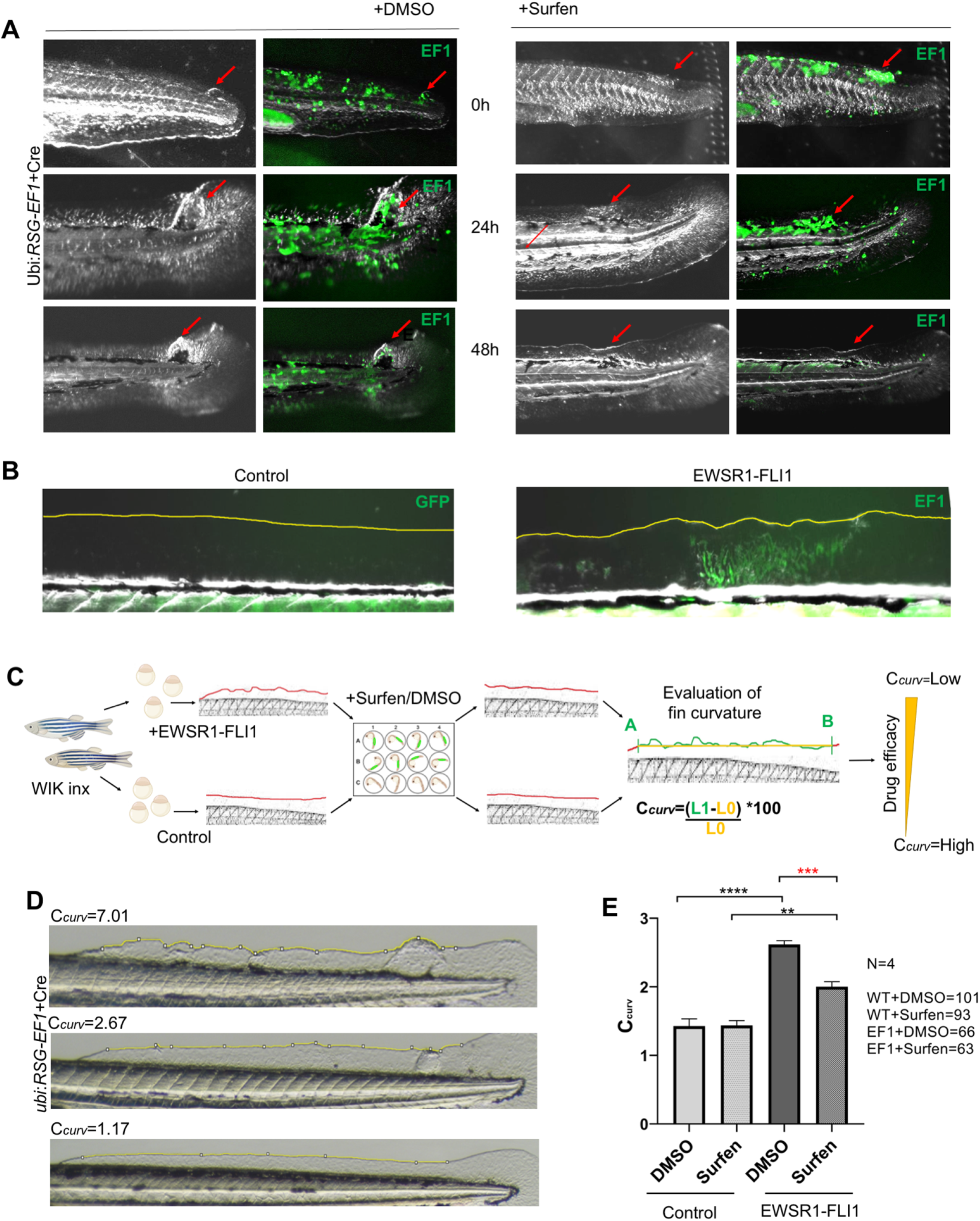
Surfen inhibits *EWSR1-FLI1* mediated growth in the zebrafish model. **A,** Surfen inhibits the development of *EWSR1-FLI1* driven outgrowths in zebrafish larvae. **B,** Changes in the fin shape driven by *EWSR1-FLI1* expression. Embryos injected with *ubi*:*RSG-EWSR1-FLI1* plus *GFP* mRNA were used as control. **C,** Scheme of surfen treatment. Wild type embryos were used for the integration of *EWSR1-FLI1* into the zebrafish genome. Uninjected fish were used as control. GFP positive embryos were treated with surfen at 2μM or 2% DMSO vehicle control. Zebrafish were imaged after 48 hours of treatment. Coefficient of curvature was calculated as following Ccurv=(L1-L0)/L0*100, where L1 is the length of fin edge between points A and B and L0 is the length of the straight line between those two points. **D,** Fins representing different coefficients of curvature Ccurv caused by *EWSR1-FLI1* expression. **E,** Treatment of fish expressing *EWSR1-FLI1* with surfen leads to the rescue of fin curvature phenotype. Error bars represent SEM, biological replicates N=4, ** indicates p<0.01, *** indicates p<0.001, **** indicates p<0.0001 based on one-way ANOVA test.

To quantify the effect of surfen, we took advantage of the changes in fin morphology driven by the EWSR1-FLI oncofusion during the first 5 days of zebrafish development. We noticed that *EWSR1-FLI1* expression is associated with distortion of the normally straight appearance of the dorsal fin border (Fig. 7B). To measure the distortion of the fin we calculated the coefficient of curvature (Ccurv), which measures the degree of deviation of the border from a straight line; more irregular fins are characterized by higher Ccurv (Fig. 7D). We hypothesized that treatment of fish with drugs targeting *EWSR1-FLI*-related pathways would result in the rescue of the fin phenotype (Fig. 7C).

Applying this metric, surfen did not affect the fin shape of control fish (Fig. 7E). *EWSR1-FLI1* expression strongly increased the Ccurv, and surfen treatment of fish expressing *EWSR1-FLI* resulted in significant rescue of the phenotype (Fig. 7E). Taken together, these results demonstrate that surfen can modulate ERK1/2 signaling and Ewing sarcoma cell growth both *in vitro* and *in vivo*.

## Discussion

Although Ewing sarcoma was first described over 100 years ago, there are few if any molecularly targeted therapies for patients with metastatic or relapsed disease. Fewer than 30% of patients presenting with metastases survive for 5 years (Riggi et al., 2021). While great progress has been made using cells, xenografts and PDX models, the development of complementary Ewing sarcoma animal models remains crucial for the understanding of disease biology in the complex developmental microenvironment. The importance of tumor microenvironment and *communication* between cancerous and host *cells has become evident as essential for tumor formation and invasion*. A better understanding of such mechanisms will support the development of new approaches, targeting not just cancer cells, but the environment around them.

Previous attempts by multiple groups to generate a mouse model of Ewing sarcoma were complicated by the high embryonic toxicity of the driver oncofusion (Minas et al., 2017). Consistent with this experience, we have tested a panel of ubiquitous and tissue specific promoters to drive *EWSR1-FLI1* expression in zebrafish embryos (Supplemental Fig.1 A, B). We found that in most cases, expression of the oncogene causes cellular apoptosis, embryonic lethality or developmental defects. Cre-inducible expression of *EWSR1-FLI* under *cmv* and *b-actin* ubiquitous promoters did not result in a high incidence of tumor development. However, we discovered that Cre-inducible expression of human *EWSR1-FLI1* under the ubiquitin promoter was associated with less oncofusion toxicity in zebrafish larva. The pattern of Cre-induced *eGFP-2A-EWSR1-FLI* expression was consistently different from Cre-induced GFP expression driven by the same ubiquitin promoter, suggesting that *EWSR1-FLI* can be tolerated by certain cell types while being toxic for others.

Human Ewing sarcoma can occur in any part of the body in bone or soft tissue, but not endodermally or ectodermally derived tissue. It most commonly involves the pelvis and proximal long bones (Riggi et al., 2021). In our model, based on mosaic integration of human *EWSR1-FLI1* into the zebrafish genome, formation of tumors was observed in association with mesenchymally derived skeleton at the regions proximal to the base of pectoral, anal and caudal fins, and at supraneurals. 58% of fish had more than one tumor (Supplemental Fig.3 A, Table 1). Interestingly, we observed the formation of ectopic fins driven by *EWSR1-FLI1* (Supplemental Fig.3 B) suggesting that *EWSR1-FLI1* may potentially re-direct the differentiation program of transformed cells. This model presents several advances over our previously-reported zebrafish mosaic model of Ewing sarcoma (Leacock et al., 2012). In that model, on a wild type background only 0.6% of fish developed tumors during the 15 months, and tp53-deficiency was required for more penetrant tumor formation. (Leacock et al., 2012). 40% of affected fish developed leukemia-like tumors rather than sarcomas. In our new Cre-inducible mosaic model, the 34% of fish developed tumors. Only 1 fish out of 77 developed a leukemia-like SRBCT. These findings highlight the important role that the level and spatiotemporal distribution of *EWSR1-FLI1* expression play in efficient tumor generation.

To characterize tumors, we performed IHC staining for markers commonly used in diagnosis of human Ewing sarcoma. Staining of zebrafish tumors with antibodies against FLI1 showed that cells have different expression level of *EWSR1-FLI*, consistent with recent reports of cell to cell heterogeneity which affects proliferative and migratory potential of Ewing sarcoma cells (Franzetti et al., 2017). While silencing of transgene expression in individual cells may occur, the fact that *GFP* and *EWSR1-FLI1* are expressed from a single mRNA transcript, and *GFP* expression is widely retained throughout the tumor, make this possibility less likely. Staining for CD99 and PAS are widely used for Ewing sarcoma diagnosis in patients (Muhammad et al., 2012). CD99 is a cell surface transmembrane protein highly expressed in all Ewing’s sarcomas. Zebrafish tumor cells were positive for CD99. Additionally, consistent with human data zebrafish tumors were positive for PAS which stains glycogen and polysaccharides enriched in Ewing sarcoma. We showed that the expression of *nr0b1* in tumor tissue is strongly upregulated resembling the human phenotype (Kinsey et al., 2006). Altogether, zebrafish tumors are positive for known Ewing sarcoma markers recapitulating the main features of the human disease.

Previous studies established that 91% of human tumors had proliferating populations of cells positive for ki67 marker. High proliferation index was predictive of poor overall survival independent of tumor site, tumor volume or metastasis at diagnosis (Brownhill et al., 2014). We identified increased proliferation in *EWSR1-FLI1* driven outgrowths in the earliest stages of tumor formation. In a wide variety of cancers, the ERK1/2 signaling pathway is known to control cell proliferation. Moreover it was shown that ERK1 and ERK2 are constitutively activated in NIH 3T3 cells expressing *EWSR1-FLI1* as well as in several human Ewing’s sarcoma tumor-derived cell lines (Silvany et al., 2000). Consistent with these data, we identified activated phosphorylation of ERK1/2 in both *EWSR1-FLI1* expressing outgrowths and tumors in the *in vivo* zebrafish model. In summary, tumorigenesis in fish is associated with activation of ERK1/2 signaling.

To study the mechanisms underlying the activation of ERK1/2 triggered by *EWSR1-FLI1* we performed LC-MS/MS analysis. We discovered that that *EWSR1-FLI1* affects expression of proteins involved in ECM reorganization and proteoglycan catabolism. It is known that tumors leverage ECM remodeling to create a microenvironment that promotes tumorigenesis and metastasis. These tumor-driven changes support tumor growth, migration and invasion. Our data show that expression of *EWSR1-FLI1* is associated with the strong production of collagens col1a1b, col1a2, col2a1a, col9a1b, col9a2, col28a2a both in embryos (Supplemental table 2) and in advanced tumors (data not shown), suggesting the importance of specific matrix for tumor development. We found a strong upregulation of proteins involved in proteoglycan catabolism in human Ewing sarcoma (Fig. 4F). Interestingly, the upregulation of heparanase, the enzyme involved in cleavage of the side chains of heparan sulfate proteoglycans, was reported in Ewing sarcoma tumors correlating with poor prognosis in patients (Shafat et al., 2011). Thus, Ewing sarcoma oncogenesis is associated with dysregulation of proteoglycan turnover.

Proteoglycans are important components of extracellular matrix regulating Wnt, Hedgehog, TGF-β, FGFR and other signaling pathways. Proteoglycans stabilize ligand–receptor interactions, creating potentiated ternary signaling complexes that regulate the signaling pathways involved in cell proliferation, migration, adhesion and growth factor sensitivity (Elfenbein and Simons, 2010). For example, binding of FGF to its signaling receptor requires prior binding to the heparan sulfate side chains of the proteoglycans (Mythreye and Blobe, 2009). Thus, dysregulation of proteoglycan catabolism can result in aberrations in signal transduction from cell surface receptors.

To block the aberrantly activated proteoglycan-mediated pathways in Ewing sarcoma we targeted the binding of GAG chains with the ligands and signal molecules. Surfen (*bis*-2-methyl-4-amino-quinolyl-6-carbamide) binds to heparan sulfate and other GAGs blocking the sulfation and degradation of GAG chains *in vitro* (Schuksz et al., 2008). Surfen also affects responses dependent on heparan sulfate such as growth factor binding and activation of downstream signaling pathways (Schuksz et al., 2008). To target proteoglycan-mediated activation of ERK1/2 signaling we treated TC32 and EWS502 cells with surfen. Surfen impaired Ewing sarcoma cell proliferation at all doses tested, with the strongest effects at 5 and 10 μM. Concomitant with the effect on proliferation, ERK1/2 phosphorylation was downregulated at these doses of surfen, indicating that Ewing sarcoma cell lines are very sensitive to surfen-mediated blockage of cell surface receptor signaling.

To test whether surfen is effective *in vivo* in the zebrafish model we treated fish with outgrowths via aqueous exposure to surfen at 2 μM. Surfen demonstrated remarkable inhibition of outgrowth development in zebrafish larvae. Thus, surfen is effective against human Ewing sarcoma cells *in vitro* and against tumor growth in the zebrafish model *in vivo*. Surfen was originally developed for use in humans to facilitate depot insulin deposition (Umber et al., 1938), and has been tested as an agent against glioblastoma tumor cells (Logun et al., 2019). Thus, targeting glycosaminoglycan metabolism may represent a new therapeutic opportunity for Ewing sarcoma. We quantified the effect of surfen *in vivo* using the fin coefficient of curvature Ccurv. We note that this assay has great potential for high-throughput screening to identify additional drugs targeting proteoglycan-mediated signaling in Ewing sarcoma.

Overall, here we present a new inducible zebrafish model of Ewing sarcoma. The advantage of the model is the opportunity to study tumorigenesis *in vivo*, in a complex developmental background. The system further allows study of tumor cell behavior using high-resolution imaging and is effective for high-throughput drug screening. Our findings suggest that the temporal expression of *EWSR1-FLI1* is crucial for tumor development, supporting the existence of a specific cell or lineage of origin for Ewing sarcoma. Our new *EWSR1-FLI* zebrafish model of Ewing sarcoma emphasizes the role of proteoglycans mediating ERK1/2 signaling and growth of tumor cells. Further investigation of the interactions between Ewing sarcoma and the tumor microenvironment *in vivo* can provide critical insights that may lead to new therapies of the disease.

## ACKNOWLEDGMENTS

This project is funded by grant U54 CA231649-01-A1 from the National Cancer Institute and by grants from the 1 Million 4 Anna Foundation and Curing Kids Cancer. We thank the Children’s Hospital Los Angeles Pathology and Cellular Imaging Cores, the UT Southwestern Medical Center Proteomics Core, and the University of Southern California High-Performance Computing Cluster for exceptional services and for their expertise. We grateful to Genevieve Kendall for productive discussions and help. JFA was previously supported by the Nearburg Professorship of Pediatric Oncology Research at UT Southwestern.

## Methods

### Zebrafish Husbandry

*Danio rerio* were maintained according to industry standards in an AALAAC-accredited facility. WIK wild type fish were obtained from the ZIRC (Zebrafish International Resource Center (https://zebrafish.org).

### Plasmids and Cloning

Human *EWSR1-FLI1* coding sequence was provided by Chris Denny, University of California-Los Angeles, USA (Leacock et al., 2012). The Gateway expression system (Invitrogen) was used for generation of all constructs for expression in zebrafish (Kwan et al., 2007). *EWSR1-FLI1* flanked by attB2r site (at 5’primer) and attB3 site (at 3’primer) was cloned into a 3’entry vector according to the provided protocol (Kendall and Amatruda, 2016). The plasmids containing a stop-dsRed-stop sequence were a generous gift from Eric Olson. Likewise dsRed-STOP-GFP-2A-*EWSR1-FLI1* coding sequence flanked by attB1 and attB2 sites was cloned into a middle entry vector (Kwan et al., 2007). The *ubi* promoter was a kind gift from Len Zon (Addgene #27320) (Mosimann et al., 2011). The *fli1* promoter was provided by Nathan Lawson (Addgene #31160 and #26031), and the *mitfa* promoter by James Lister (Addgene #81234) (Kendall et al., 2018). The *beta actin* promoter, *cmv* promoter were used for plasmids generation and expression in zebrafish. The plasmids containing a GFP-2A sequence were a kind gift from Steven Leach, and were sub-cloned into a middle entry Gateway expression system (Kendall et al., 2018). The destination vector pDestTol2pA2 and 3’ SV40 late poly A signal construct, were used for construct generation by an LR reaction with LR Clonase II Plus (Invitrogen) (Kwan et al., 2007). Transposase, GFP and Cre RNAs were synthesized from plasmids pCS2FA, pCS2-Cre.zf and pCS2-GFP accordingly using the mMessage mMachine kit (Applied Biosystems/Ambion, Foster City, CA). The constructs generated include: *beta-actin-GFP2A-pA* (Genevieve Kendall), *beta-actin-GFP2A-EWSR1-FLI1*, *ubi-eGFP-2A*, *ubi-GFP2A-EWSR1-FLI1*, *cmv-eGFP-2A*, *cmv-GFP2A-EWSR1-FLI1*, *fli-eGFP-2A*, *FLI-GFP2A-EWSR1-FLI1*, *mitfa-eGFP-2A*, *mitfa-GFP2A-EWSR1-FLI1*, *beta-actin-dsRed-stop-eGFP-2A-pA*, *beta-actin-dsRed-stop-eGFP-2A-EWSR1-FLI1*, *ubi-dsRed-stop-eGFP-2A*, *ubi-dsRed-stop-eGFP-2A-EWSR1-FLI1*, *cmv-dsRed-stop-eGFP-2A*, *cmv-dsRed-stop-eGFP-2A-EWSR1-FLI1*, *fli-dsRed-stop-eGFP-2A*, *fli-dsRed-stop-eGFP-2A-EWSR1-FLI1*, *mitfa-dsRed-stop-eGFP-2A*, *mitfa-dsRed-stop-eGFP-2A-EWSR1-FLI1*.

### Zebrafish injections

Zebrafish embryos were injected at the single-cell stage. The injection mixes contained 50 ng/μL of Tol2 transposase mRNA, 10-50 ng/μL of described DNA constructs, 0.1% phenol red, and 0.3X Danieau’s buffer. The injection mixes containing *beta-actin-dsRed-stop-eGFP-2A-pA*, *beta-actin - dsRed-stop-eGFP-2A-EWSR1-FLI1*, *ubi-dsRed-stop-eGFP-2A*, *ubi-dsRed-stop-eGFP-2A-EWSR1-FLI1*, *cmv-dsRed-stop-eGFP-2A*, *cmv-dsRed-stop-eGFP-2A-EWSR1-FLI1*, *fli-dsRed-stop-eGFP-2A*, *fli-dsRed-stop-eGFP-2A-EWSR1-FLI1*, *mitfa-dsRed-stop-eGFP-2A*, *mitfa-dsRed-stop-eGFP-2A-EWSR1-*FLI1 also contained 0.5 ng of *Cre*_RNA or *GFP*_RNA.

### Zebrafish embryo survival

For survival analysis of embryos, fish were injected with *ubi*:*RSG-EWSR1-FLI1; Cre*_RNA or control mixes *ubi*:*RSG-EWSR1-FLI*; *GFP*_RNA *ubi*:*RSG; Cre*_RNA. The total number of injected fish was counted (the exact number of fish for each condition is indicated in the respective legend), and then the resulting alive embryos subsequently determined during first 10 days. Survival curves were plotted using GraphPad Prism 8.4.3. Biological replicates N= 3.

### Protein Identification by LC-tandem Mass Spectrometry

Zebrafish embryos were injected with *ubi*:*RSG-EWSR1-FLI1;Cre*_RNA or *ubi*:*RSG;Cre_RNA*. Embryos were sorted for GFP at 24hpf and 48hpf time points. Sorted embryos were dechorionated, dissociated by pestle homogenizer in RIPA buffer supplemented with 1x inhibitor of proteases (Mini Protease Inhibitor Cocktail, cOmplete). Additionally, tumor or normal tissue from dorsal, caudal and pectoral fins were dissected and processed in an analogous way. Protein levels were analyzed using the BCA protein assay Kit (Thermo Scientific). Lysates were heated at 95°C for 5 minutes and loaded on a 12% gel (BioRad). Gels were stained with Coomassie Blue. Stained 1cm bands were cut out of gels, sliced into 1mm cubes and transferred to 1.5 Eppendorf tubes for submission. Data were pre-proceeded and provided by the UT Southwestern Proteomics Core. Data were analyzed using an R package ROTS (Suomi et al., 2017) and visualized in R. Gene Set enrichment analysis was performed using the GSEA software according the instructions provided https://www.gsea-msigdb.org/gsea. The experiment was performed in 3 technical replicates for embryo samples and 3 biological replicates for tissue samples.

### Zebrafish tumor collection, processing for histology, and tumor incidence

Zebrafish were screened under a Nikon SMZ25 fluorescent stereomicroscope for the presence of GFP positive tumors. Fish with tumors were humanely euthanized and fixed in 4% paraformaldehyde/1XPBS for 48 hrs at 4°C. They were de-calcified in 0.5M EDTA for 5 days, processed and mounted in paraffin blocks for sectioning and further experiments. For tumor incidence curves, zebrafish were injected and sorted for GFP at 14 dpf and monitored for 4 months. Zebrafish with no GFP fluorescence were considered as negative for *EWSR1-FLI-* dependent tumor formation. Zebrafish with tumors were collected and processed as described earlier for hematoxylin and eosin staining. All tumors were reviewed by an experienced sarcoma pathologist.

### Immunofluorescence

In zebrafish embryo whole mounts, embryos at 72hpf were fixed overnight at 4 °C in 4% PFA in 1xPBS and processed according to a published protocol (Verduzco and Amatruda, 2011). Zebrafish embryos were stained with primary antibodies directed against Phospho-p44/42 MAPK (Erk1/2) Thr202/Tyr204 (4370S, Cell Signaling) at 1:200, GFP (4B10) (2955, Cell Signaling) at 1:300, or Phospho-Histone H3 (Ser10) (D7N8E) (53348, Cell Signaling) at 1:500. The secondary antibodies used were Goat anti-Mouse IgG (H+L) Alexa Fluor Plus 488 (# A32723, Thermo Fisher), Donkey anti-Rabbit IgG (H+L) Alexa Fluor 546 (A10040, Thermo Fisher), Goat anti-Rabbit IgG(H+L) Alexa Fluor 405 (A31556, Thermo Fisher) at 1:500. The staining was repeated at least 3 times.

### IHC staining

Slides with paraffin embedded tissue sections were baked for 60 minutes at 60°C, immersed with xylene, 100% ethanol, 95% ethanol, 75% ethanol, distilled H2O two times each for 5 minutes each. Antigen retrieval was performed in Trilogy reagent (920P, Sigma) for 10 minutes in the pressure cooker. Slides were cooled and blocked with 3% H2O2 for 30 minutes, followed by blocking with 1%BSA/1xPBST for 1 h. Slides were incubated with primary antibodies Anti-CD99 antibody (ab108297, Abcam) at 1:200, Anti-FLI1 (ab15289, Abcam) at 1:100, anti-Phospho-p44/42 MAPK (Erk1/2) Thr202/Tyr204 (4370S, Cell Signaling) at 1:200 overnight. Secondary antibodies used were Anti-rabbit IgG, HRP-linked Antibody (7074S, Cell Signaling), Anti-mouse IgG, HRP-linked Antibody (7076S, Cell Signaling). SignalStain^®^ DAB Substrate Kit #8059 was used for chromogen staining according to manufacturer’s instructions. Slides were also stained with hematoxylin and eosin, dehydrated and mounted with permount mounting media. The staining was repeated more than 3 times.

### Imaging

Embryos were imaged at 12 hpf, 24 hpf, 48 hpf, 72 hpf, and 5 dpf on Nikon SMZ25 fluorescent stereomicroscope. Images of whole mount immune-stained zebrafish were taken on a Keyence BZ-X700 fluorescent microscope. Slides were imaged on a Keyence BZ-X700 fluorescent microscope, Leica DM4000B and Zeiss LSN710.

### RNA extraction, cDNA synthesis, and qRT-PCR

Fresh GFP positive tumor tissues or sorted dechorionated embryos were snap frozen in liquid nitrogen. Frozen tissues and embryos were subjected to total RNA isolation using the RNeasy Microkit manufacturer’s instructions (Qiagen). The RT2 HT First Strand Synthesis kit (QIAGEN) was used for cDNA reverse transcription from 200 ng-1μg of total RNA. qRT-PCR was performed on a CFX384 Touch Real-Time PCR Detection System using the SYBRGreen Master Mix (BioRad). See <u>Supplemental Table 3</u> for primer sequences. Error bars indicate standard deviation. To determine significance a student’s t-test was performed on normalized Ct replicates. The RT-PCR analysis was performed at least 3 times for each experiment in 3 technical replicates for each condition.

### Cell culture

TC32 and EWS502 cell lines were the kind gift of Dr. Angelique Whitehurst. TC32 was maintained in RPMI 1640 GlutaMAX with 10% Fetal Bovine Serum (Sigma) and 1X Antibiotic-Antimycotic (Gibco) at 37°C in 5% CO_2_. EWS502 was maintained in RPMI 1640 GlutaMAX with 15% FBS (Sigma) and 1X Antibiotic-Antimycotic (Gibco) at 37°C in 5% CO_2_.

### Surfen treatment

Surfen hydrate, ≥98% was obtained from Sigma-Aldrich (S6951). Because Surfen binds avidly to plastic, it is necessary to use glass vessels or precoat all plasticware with serum before use. Surfen stock solutions were prepared in DMSO at 1 mM and 5 mM, aliquoted and stored in glass containers at −20°C in the dark.

### Zebrafish treatment with surfen

For zebrafish treatment working solutions were prepared by diluting stock solutions in E3 medium. Final concentrations for treatments were determined as 2 μM for surfen hydrate (S6951, Sigma-Aldrich), based on maximal non-toxic concentrations for zebrafish embryos (Naini and Soussi-Yanicostas, 2018). Control fish were treated with 0.2% DMSO. Embryos were injected, screened for GFP at 24 hpf, dechorionated and incubated for 2 days in 10 ml of either control medium (E3 medium containing 0.2% DMSO) or E3 medium with 2mM surfen in a glass beaker. All solutions were changed daily. Biological replicates N= 4. The exact number of fish for each condition is indicated on figure legend.

### Protein extraction and western blots

Cells or tumor tissues were harvested and lysed in RIPA buffer complemented with cOmplete Mini Protease Inhibitor Cocktail inhibitors and Phosphatase inhibitor cocktails (Sigma). Lysates were denatured by boiling in 1x Laemmli buffer at 95°C for 5 minutes and loaded on a 4–20% gradient gel (BioRad). PVDF membranes were used for proteins wet transfer (Bio-Rad). Membranes were blocked in Casein + 0.1% Tween-20 (Thermo), and incubated overnight at 4°C with the primary antibodies Phospho-p44/42 MAPK (Erk1/2) (Thr202/Tyr204) (4370S, Cell Signaling) at 1:2000, p44/42 MAPK (Erk1/2) (137F5) (4695S, Cell signaling) at 1:2000, β-Actin Antibody (4967S, Cell Signaling) at 1:2000, α-Tubulin Antibody (2144S, Cell Signaling) at 1:2000. Secondary antibody used were Anti-rabbit IgG, HRP-linked Antibody (7074S, Cell Signaling) at 1:5000, Anti-mouse IgG, HRP-linked Antibody (7076S, Cell Signaling) at 1:5000. Signal was detected using SuperSignal West Pico Chemiluminescent Substrate (Fisher). BioRad GelDoc XR was used for membranes imaging. The western blot analysis was performed at least 3 times for each experiment in 3 technical replicates.

### Cellular proliferation assays

For proliferation TC32 and EWS502 were seeded at 2,000 cells per well of 96-well plate with three replicas per timepoint in 10% and 15% RPMI 1640 GlutaMAX media supplemented with 1X Antibiotic-Antimycotic (Gibco). Cells were treated with 1.75uM, 2.5uM, 5uM and 10 uM of Surfen or 0.025%, 0.05%, 0.1%, 0.2% DMSO on 2nd, 4th, 6th, 8th days. The number of cells at each point was measured using the hemocytometer at 1st, 3rd, 5th, 7th, 9th days. The proliferation assay was performed at least 2 times. Three technical replicates were performed per condition.

### Colony formation assay

For colony formation assay 500 of TC32 and EWS502 cells were seeded per well of 24 wells low adhesive plate treated with 0.5 uM, 1uM, 1.5um, 2uM, and 2.5uM of surfen or 1% DMSO under serum-deprived conditions. Serum was added 12 hours later. Medium with surfen or DMSO was changed 3 times a week. The colony number was analyzed on day 14 and 21 for TC32 and EWS502 accordingly. For imaging media was removed, cells were pre-fixed and stained with 0.5% crustal violet. The colony formation assay was performed at least 2 times. Four technical replicates were performed per condition.

### Ccurvature calculation

To measure the distortion of the fin we introduced the coefficient of curvature C*curv*=(L1-L0)/L0*100, where L1 is the length of the fin border between points A and B and L0 is the length of the straight line between those two points (Fig7 C).

### Statistics

Statistical analysis for embryonic survival, tumor incidence curves, proliferation curves, colony formation, relative gene expression, were performed using GraphPad Prism 8.4.3 (La Jolla, CA). The number of individual experiments, replicas and samples analyzed, and significance are reported in the figure legends. Statistical significance was calculated by Student’s t-test for two-group comparison, one-way ANOVA for comparison of multiple groups with one control group and for comparison between different experimental groups. p>0.05 = n.s, p<0.05 = *, p<0.01 = **, p<0.001 = *** and p<0.0001 = ****.

## Supplemental Materials

### Supplemental Figure Legends

**Supplemental Figure 1, referring to Figure 1.**
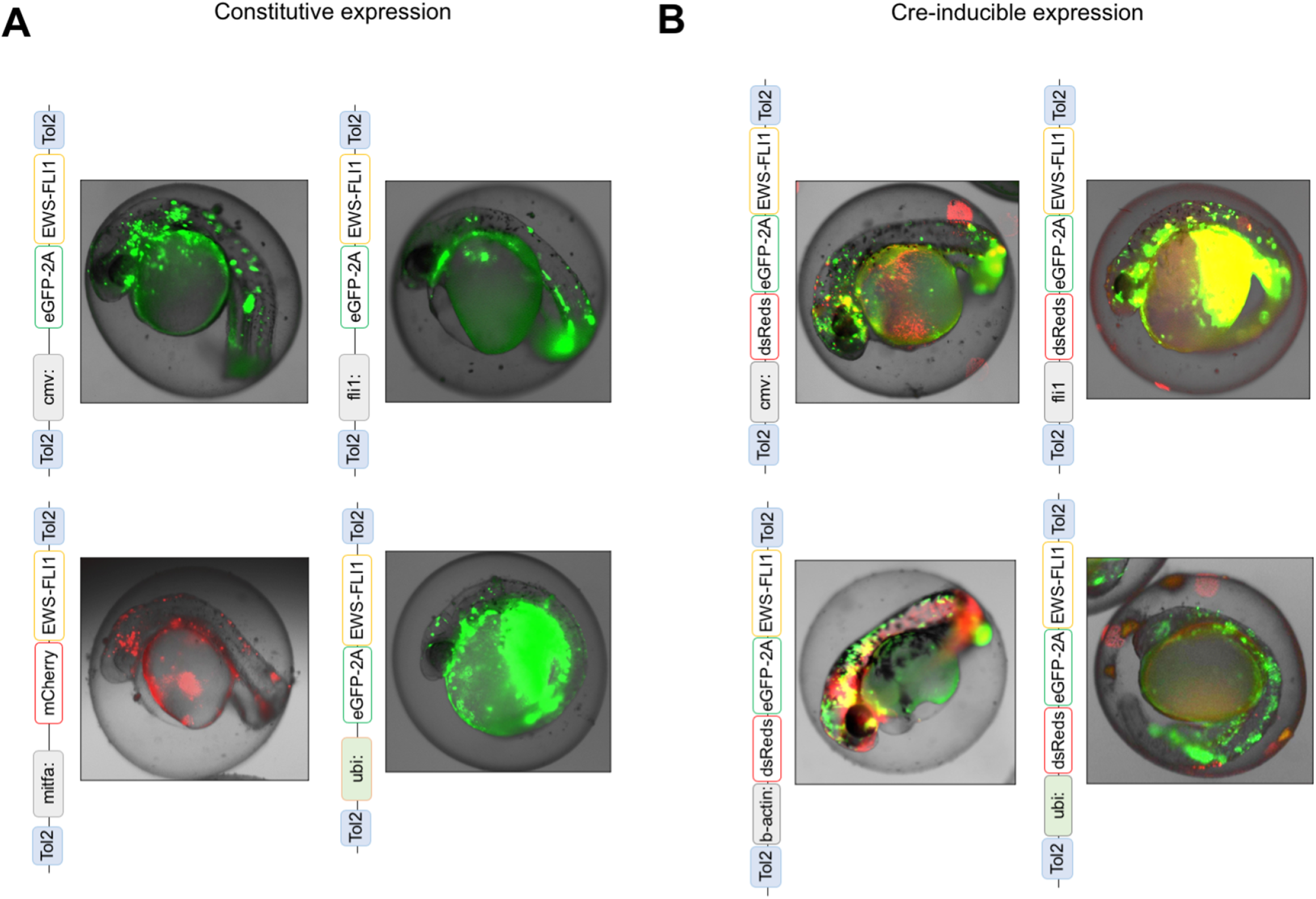
The Tol2 transposon-based system was used to integrate *EWSR1-FLI1* into the zebrafish genome by microinjection into single-cell stage zebrafish embryos. **A,** Scheme of constructs for expression of *EWSR1-FLI1* under *cmv*, *fli1*, *mitfa*, *ubi* promoters. **B,** Scheme of constructs for Cre-inducible expression of eGFP-2A-*EWSR1-FLI1* under *cmv*, *fli1*, *b-actin*, *ubi* promoters.

**Supplemental Figure 2, Referring to Fgure 1.**
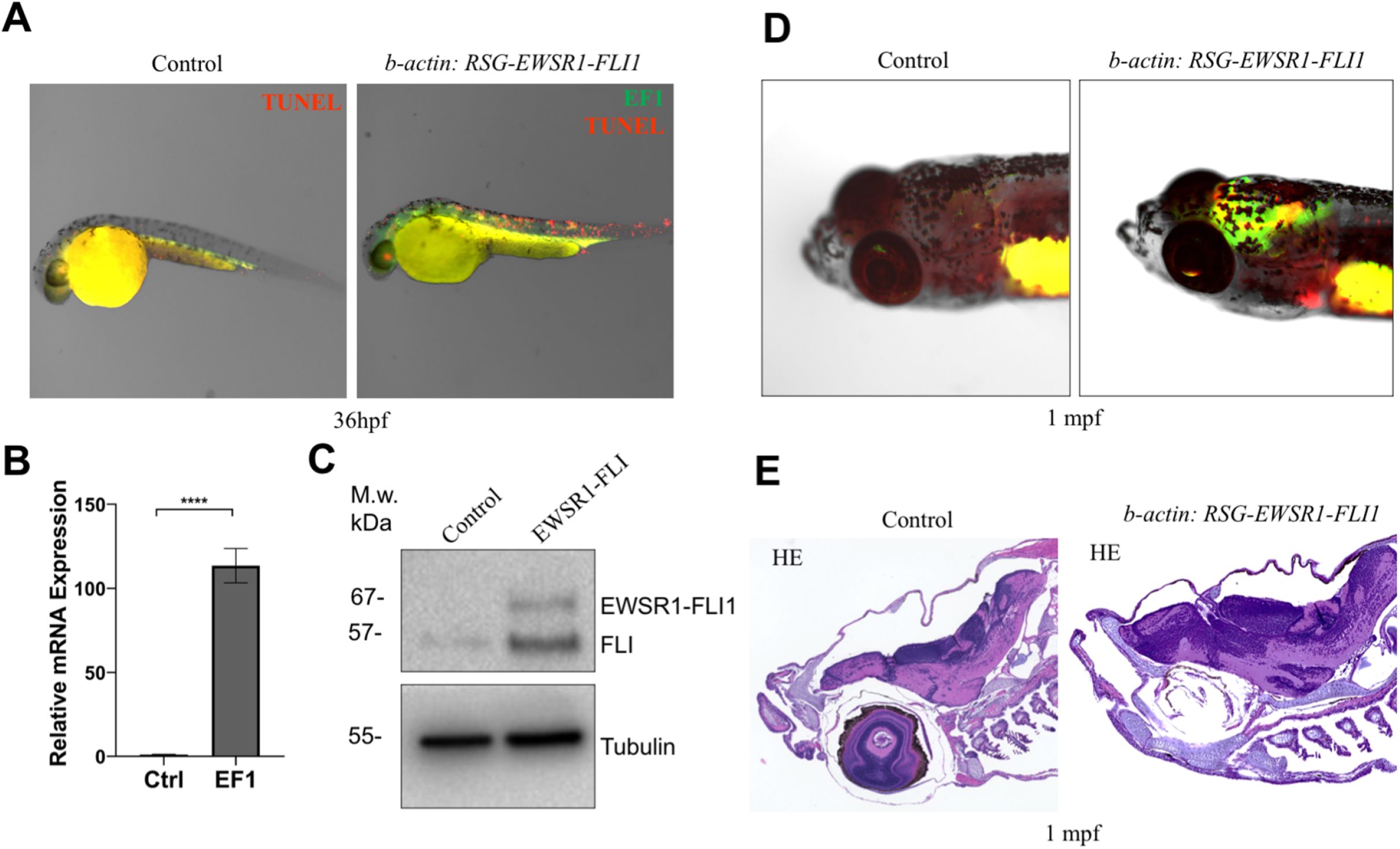
The Tol2 transposon-based system was used to integrate *b-actin:RSG-EWSR1-FLI1* into the wild type zebrafish genome. Coinjection of *Cre* mRNA or *GFP* mRNA was used to generate *EWSR1-FLI1* expressing or control zebrafish, respectively. **A,** TUNEL assay (red) made on zebrafish expressing *GFP* (green) or *GFP2A-EWSR1-FLI1* (green) at 36hpf. **B,** Relative mRNA expression of *EWSR1-FLI1* in *b-actin:RSG-EWSR1-FLI1* zebrafish injected with *Cre* RNA or *GFP* RNA. **C,** Expression of EWSR1-FLI1 on the protein level wa*s* confirmed by immunoblotting*. **D,*** Representative images of zebrafish expressing *EWSR1-FLI1* driven by *b-actin* promoter at 1mpf. **E,** H&E staining of cell masses formed in the head of zebrafish expressing *EWSR1-FLI1* under *b-actin* promoter (right) and control (left). Error bars represent SEM, N = 3, **** indicates p<0.0001, two-tailed Student’s t-test.

**Supplemental Figure 3, Referring to Figure 1.**
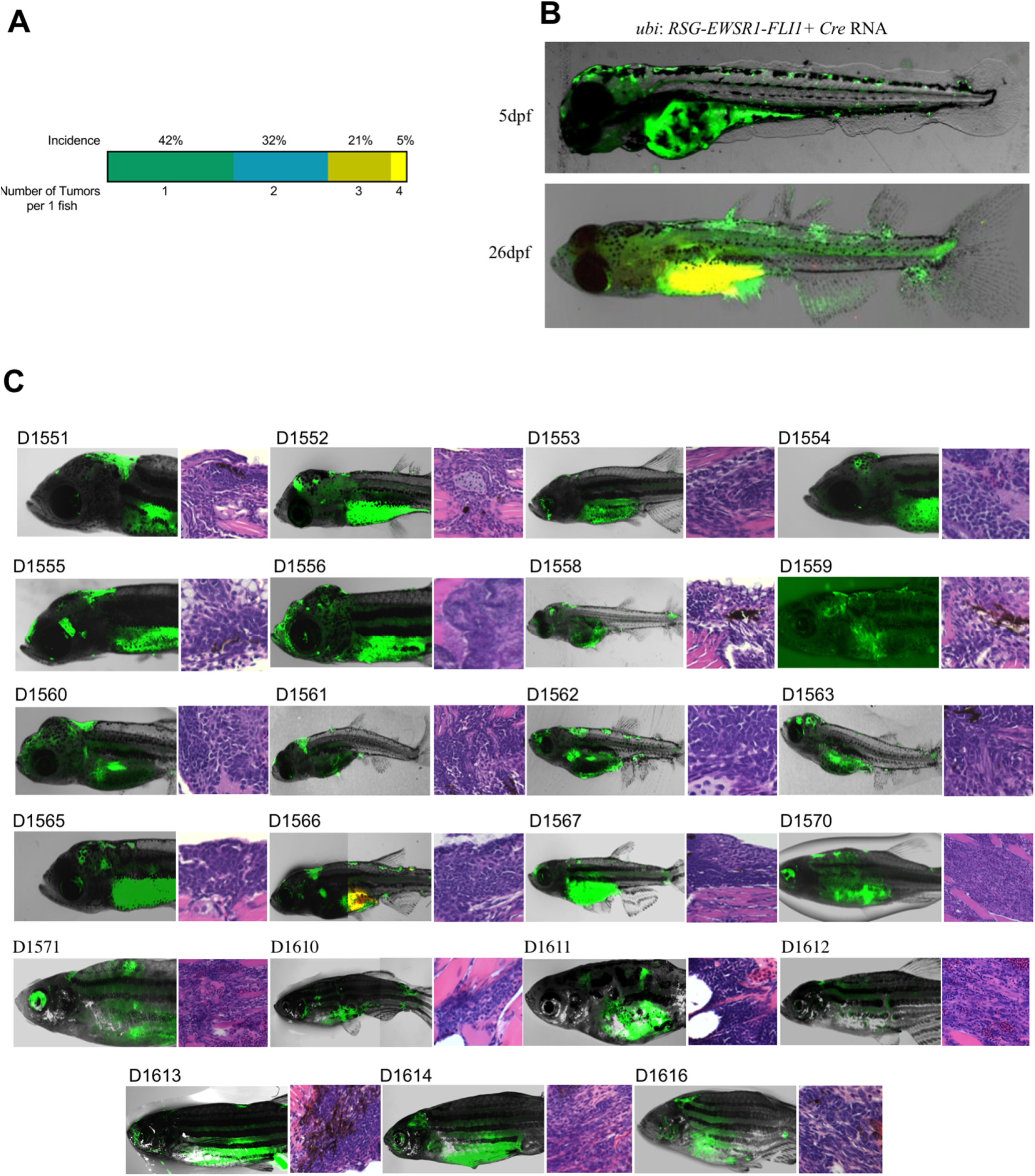
Mosaic model of Ewing sarcoma generated by the microinjection of *ubi*:*RSG-EWSR1-FLI1* plus *Cre* mRNA into single-cell stage embryos. **A,** Plot representing the incidence of one, two, three or four tumors per zebrafish. **B,** *EWSR1-FLI1* triggers the formation of ectopic fins in zebrafish. **C,** Examples of tumors observed in mosaic model of Ewing sarcoma.

**Supplemental Figure 4, Referring to Figure 2.**
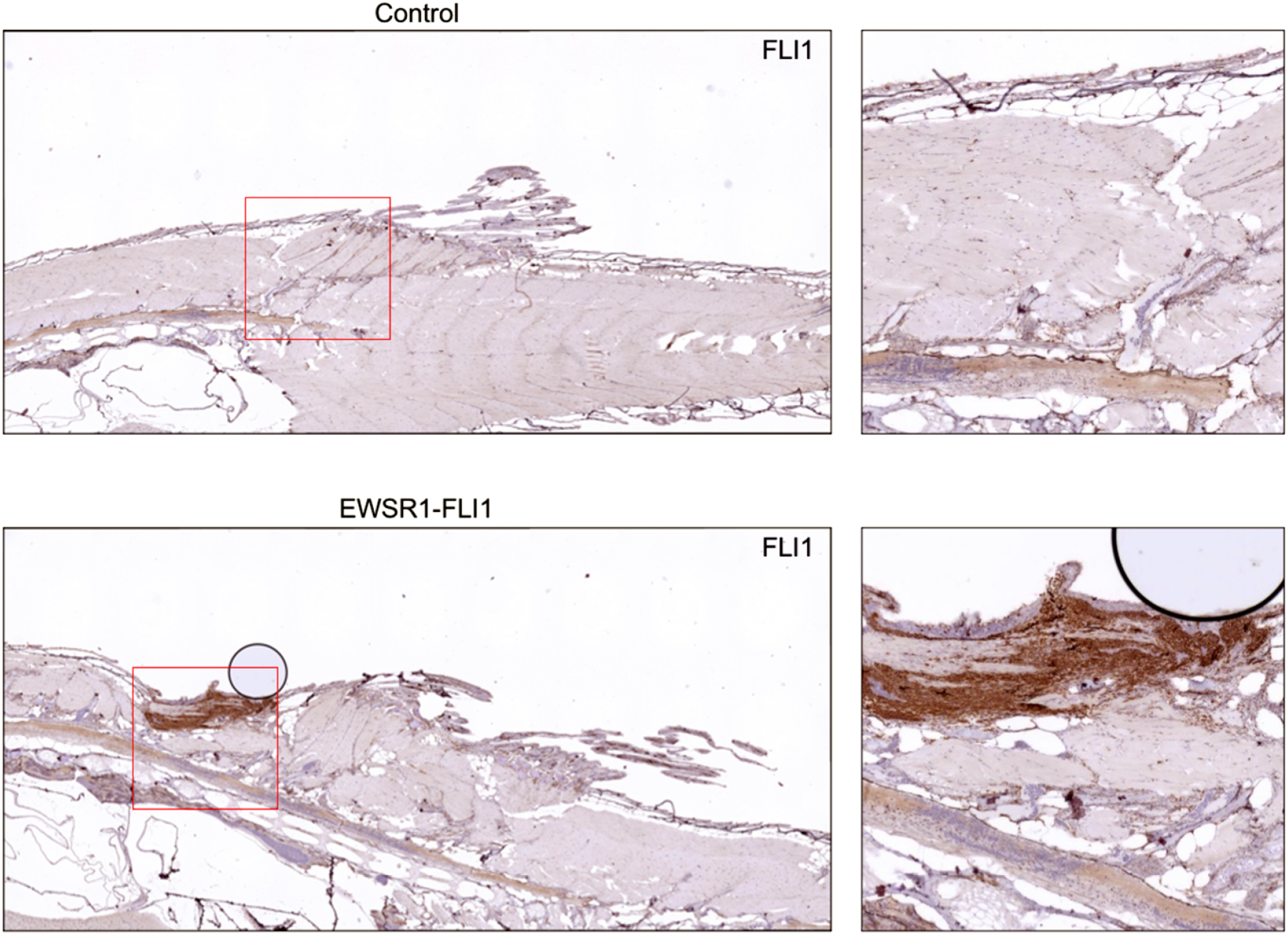
IHC staining of sections of zebrafish tumor and control zebrafish with anti-FLI1 antibodies.

**Supplemental Figure 5, Referring to Figure 4.**
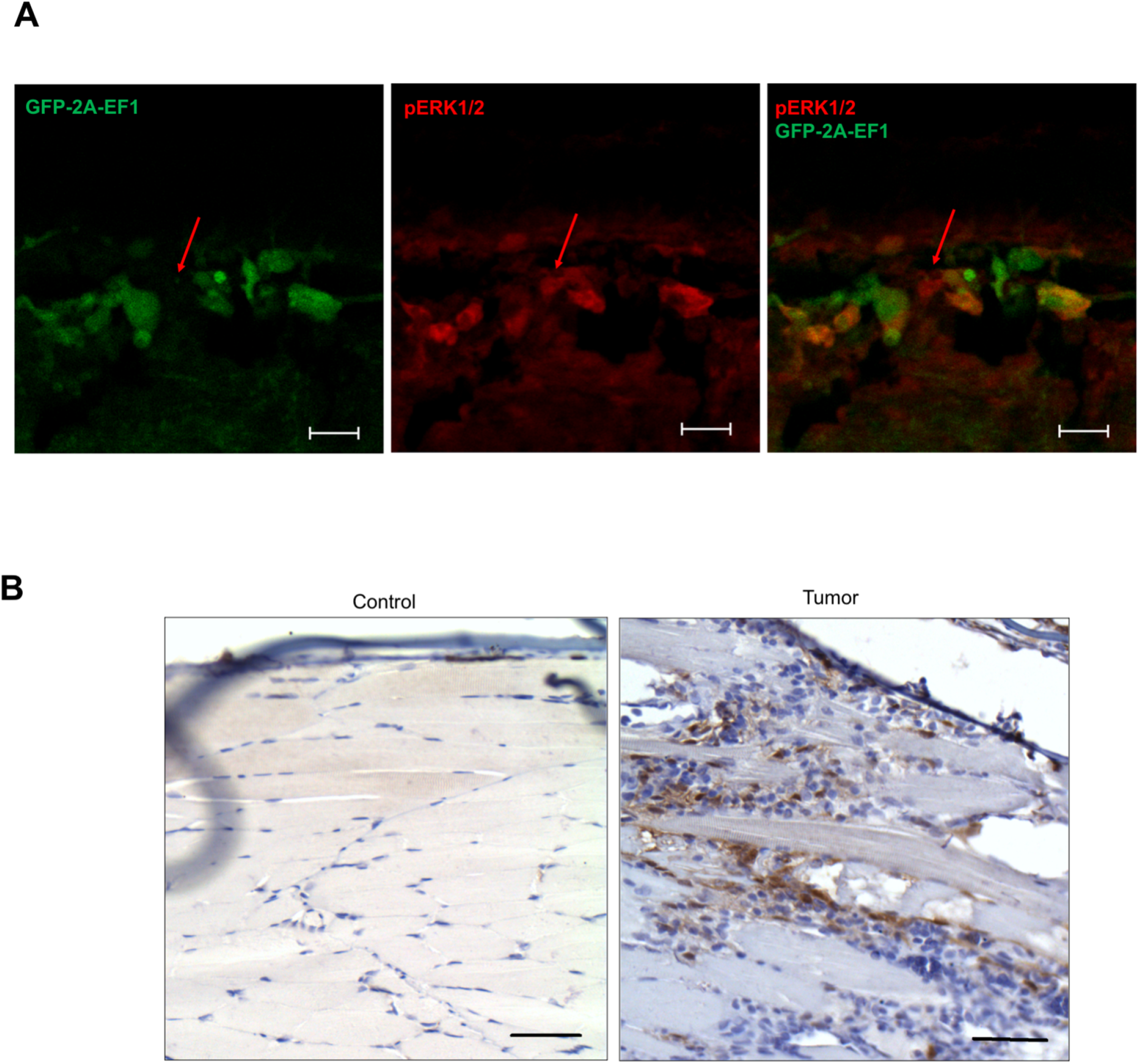
**A,** Immunofluorescence staining of zebrafish outgrowth at 72hpf for pERK1/2 (red), GFP2A-EWSR1-FLI1 (green). Arrows indicate cells positive for pERK1/2 and negative for EWSR1-FLI1. Scale bars, 20μm. **B,** IHC staining of sections of zebrafish tumor and control zebrafish with anti-pERK1/2 antibodies. Scale bars, 100 μm

**Supplemental Movie 1**

Immunofluorescence staining of a 48hpf zebrafish embryo for pERK1/2 (red), EWSR1-FLI1 (green) and pH3 (blue). Confocal Z-stack focusing on a region of the trunk and dorsal fin.

**Supplemental Movie 2**

Immunofluorescence staining of a tumor outgrowth in 48hpf zebrafish embryo for pERK1/2 (red), EWSR1-FLI1 (green) and pH3 (blue). Confocal Z-stack focusing on an outgrowth arising from the dorsal surface of the embryo.

**Supplemental Table 1**

Detailed characterization of zebrafish tumors driven by *EWSR1-FLI1* expression

**Supplemental Table 2**

List of downregulated and upregulated proteins identified by LC-MS/MS analysis *in EWSR1-FLI1* expressing embryos, *p* < 0.05

**Supplemental Table 3**

List of primers for RT-PCR

